# AHL10 phosphorylation determines RRP6L1 chromatin association and growth suppression during water stress

**DOI:** 10.1101/2023.04.17.537135

**Authors:** Min May Wong, Xin-Jie Huang, Yu-Chiuan Bau, Paul E. Verslues

## Abstract

Phosphorylation of AHL10, one of the AT-hook family of plant-specific DNA binding proteins, is critical for growth suppression during moderate severity drought (low water potential, ψ_w_) stress. To understand how AHL10 phosphorylation determines drought response, we identified putative AHL10 interacting proteins and further characterized interaction with RRP6L1, a protein involved in epigenetic regulation. RRP6L1 and AHL10 mutants, as well as *ahl10-1rrp6l1-2*, had similar phenotype of increased growth maintenance during low ψ_w_. Chromatin precipitation demonstrated that RRP6L1 chromatin association increased during low ψ_w_ stress and was dependent upon AHL10 phosphorylation. Transcriptome analyses showed that AHL10 and RRP6L1 have concordant effects on expression of stress- and development-related genes. Together these results indicate that stress signaling can act via AHL10 phosphorylation to control the chromatin association of the key regulatory protein RRP6L1. AHL10 and RRP6L1 interaction in meristem cells is part of a mechanism to down-regulate growth during low ψ_w_ stress. Interestingly, loss of AHL13, which is homologous to AHL10 and phosphorylated at similar C-terminal site, blocked the enhanced growth maintenance of *ahl10-1*. Thus, AHL10 and AHL13, despite their close homology, are not redundant but rather have distinct roles, likely related to the formation of AHL hetero-complexes.

**Summary Statement:** Phosphorylation of *Arabidopsis thaliana* AHL10 is important to control growth during drought stress, a function which is distinct from the closely related AHL13. One way that phosphorylation determines AHL10 function is by altering its ability to mediate chromatin recruitment of RRP6L1.

## Introduction

Protein phosphorylation and de-phosphorylation is a central feature of signaling pathways that control plant responses to environmental stimuli. Members of the Clade A Type 2C Protein Phosphatases (PP2Cs) have established roles in stress signaling (Cutler et al., 2010; Danquah et al., 2014; Bhaskara et al., 2019). However, there is diversity of physiological function among the Clade A PP2Cs despite their close phylogenetic relatedness. HAI1, in particular, has relatively weak effect on ABA sensitivity of seed germination, the canonical phenotype used to define the ABA signaling function of other Clade A PP2Cs, but has strong effect on growth and osmotic adjustment during low water potential (low ψ_w_) stress (Bhaskara et al., 2012; Wong et al., 2019). At the molecular level, it has also been shown that HAI1, but not the Clade A PP2Cs ABI1 or ABI2, could directly inactivate MPK3 and MPK6 (Mine et al., 2017).

Phosphoproteomic analysis of *hai1-2* identified putative HAI1-regulated phosphorylation sites on nuclear-localized proteins (Wong et al., 2019), consistent with the predominant nuclear localization of HAI1 (Antoni et al., 2011; Bhaskara et al., 2012). These included S313 and S314 of AT-Hook Like10 (AHL10). These AHL10 phosphorylation sites had been observed in a number of previous phosphoproteome profiling studies but the functional significance of AHL10 S313 and S314 phosphorylation was not known. Further experiments demonstrated that HAI1 could directly dephosphorylate the AHL10 C-terminal region containing S313 and S314 (Wong et al., 2019). Analysis of *hai1-2* and *35S:HAI1* plants indicated that HAI1 acts to attenuate low ψ_w_-induced AHL10 phosphorylation. Further characterization of AHL10 S313 and S314 phosphomimic and null mutations found that these phosphorylation sites are important for controlling growth during low ψ_w_ and also important in determining whether AHL10 forms nuclear foci or is more dispersed in the nucleoplasm (Wong et al., 2019).

The phosphoproteomic data of Wong et al. (2019) also contained a phosphopeptide from the equivalent site on one of the most closely related AHL10 homologs (AHL13 S376) (Wong et al., 2019). In a separate study, phosphoproteome analysis of chromatin-associated proteins found that AHL13 S376 was phosphorylated by Mitogen Activated protein kinases (MPKs) in response to the pathogen elicitor flg22 (Rayapuram et al., 2021). They found that AHL10 S314 was also a putative MPK6 phosphorylation site but was not responsive to flg22 treatment. Thus, phosphorylation of the C-terminal region of AHL10 and AHL13 may be regulated by MPKs and HAI1 (both directly and by HAI1 suppression of MPK3 and MPK6 activity); however, the two AHLs differ in how their phosphorylation status is regulated in response to specific environmental stimuli. AHL10 and AHL13 are Clade B AHLs (AHL1 to AHL14), which are less studied than the Clade A AHLs (AHL15 to AHL29; Zhao et al., 2013). The C-terminal regions of the Clade B AHLs are highly variable and the MPK-HAI1 regulated phosphorylation sites identified in AHL10 and AHL13 are not conserved among other Clade B AHLs (Wong et al., 2019).

AHL10 and AHL13 are members of the AT-Hook family of plant-specific DNA binding proteins. AHLs bind AT-rich DNA sequences via their N-terminal AT-hook domain (Fujimoto et al., 2004; Zhao et al., 2013). AHLs also contain a Plant and Prokaryote Conserved (PPC)/DUF296 domain which is involved in self-interaction, or cross-interaction among AHLs, to form trimers (Fujimoto et al., 2004; Seo and Lee, 2021). In contrast to many other types of transcriptional regulators, AHLs bind the minor groove of DNA and it has been proposed that binding of AHL complexes to AT-rich sites can modify DNA architecture to bring together loops of DNA and place otherwise distal elements in close proximity to induce the formation of transcriptional regulatory complexes (Zhao et al., 2013; Franco-Zorrilla et al., 2014).

AHLs are not thought to have transcriptional regulatory activity of their own. Rather, the highly variable AHL C-terminal domain may recruit other transcriptional regulators or may mediate interactions that determine AHL sub-nuclear localization. Varied AHL protein interactions may explain observations that even though different AHLs bind to essentially identical AT-rich sequences (Franco-Zorrilla et al., 2014), they can either induce or repress gene expression. For example, ChIP-sequencing of AHL29 found that binding motifs of TCP and bHLH transcription factors were more highly enriched than AT-rich sequences directly bound by AHL29 (Favero et al., 2020), consistent with previous data showing that AHL29 interacts with TCP transcription factors (Zhao et al., 2013). Such observations raise the question of whether AHLs have sufficient sequence specificity of their own to recognize specific genomic sites or whether it is the AHLs that are recruited by other proteins and then interact with nearby AT-rich sequences in a more non-specific manner. For, AHL10 specifically, we are aware of only one study examining its cellular function (other than the phosphoproteomic studies mentioned above). AHL10 was found to interact with Admetos (ADM), a protein involved in hybrid seed lethality after interploidy hybridization, and with the SET domain-containing SU(VAR)3–9 homolog SUVH9, a protein involved in deposition of the heterochromatin mark H3K9me2 (Jiang et al., 2017).

AHLs have also been associated with nuclear matrix attachment regions (Fujimoto et al., 2004) and can be involved in histone modifications at these sites (Lee and Seo, 2017; Morisawa et al., 2000; Xu et al., 2013; Xu et al., 2023). Nuclear matrix-associated AHLs are typically localized in foci within nucleoplasm. Similar foci were also observed for YFP-tagged AHL10 and for AHL10 homocomplexes visualized using Bi-molecular Fluorescence Complementation (BiFC). In both cases these nuclear foci were abolished by the AHL10 S314A phosphonull mutation (Wong et al., 2019). Together these results illustrate how functional diversity among AHLs is likely due to their interaction with a wide range of proteins via their variable C-terminal domains and association with many types of AT-rich regions in the genome. Although AHLs are sometimes referred to as transcription factors, there is still much unclarity about how they function in gene regulation.

To better understand why AHL10 S314 phosphorylation is so critical for its function to restrict growth during drought stress, we searched for AHL10-interacting proteins. One of those proteins, RRP6L1, was found to also be a suppressor of growth during low ψ_w_ stress. RRP6L1 (also referred to as Atrimmer1) is an RNA binding protein with several proposed functions related to small RNA production and RNA-directed DNA Methylation (RdDM) (Zhang et al., 2014, Ye et al., 2016; Wendte et al., 2017). Both AHL10 and RRP6L1 act by being recruited to specific sites in the genome; DNA binding sites in the case of AHL10 or sites where specific RNAs are being transcribed in the case of RRP6L1 (Shin and Chekanova, 2014; Zhang et al., 2014; Wendte et al., 2017). However, in both cases it is unclear how this occurs as neither protein is thought to have sequence specific binding.

Surprisingly, we found that RRP6L1 chromatin recruitment depended upon AHL10 and AHL10 S314 phosphorylation. We also made the unexpected discovery that AHL10 and AHL13 are not functionally redundant, despite their close homology and common regulation by HAI1 and MPKs.

## Methods

### Yeast Two-Hybrid Screening

Yeast two-hybrid screening was performed according to the manufacturer’s instructions using the ProQuest^TM^ Two-hybrid system (Invitrogen). *AHL10* full length coding sequence was cloned into the entry vector pENTR-D-TOPO and recombined into the bait destination vector pDEST32 with GAL4 DNA binding domain (DBD). The yeast two hybrid cDNA library was constructed from the mRNA of seedlings exposed to -1.2 MPa stress for 96 h according to the CloneMiner^TM^ II cDNA library construction kit (Invitrogen) and then cloned into prey vector pDEST22. The library used and further details of its construction have been previously described (Chong et al., 2019). Co-transformed yeast (MaV203 strain) containing both the bait and the prey constructs using the lithium acetate method were spread on the selection plate SC-Leu-Trp-His with 50 mM 3-AT which was sufficient to suppress auto-activation. A total of 20 µg cDNA was used and approximately 3.36 × 10^5^ colony forming units were screened. Large colonies grown on the selection plates were picked and re-streaked on the YPAD plates after 4-5 days at 30°C and some smaller colonies were picked after 8 days. Selected colonies were grown on the YPAD plates overnight at 30°C and interaction were confirmed by β-galactosidase filter lift assay. Confirmed colonies were then cultured, DNA isolated by yeast plasmid miniprep (Zymoprep) and sequenced. Gene sequences of AHL10 and truncated RRP6L1 used for yeast two-hybrid assays were cloned into the pENTR-D-TOPO and recombined into pDEST32 and pDEST22 for β-galactosidase filter lift assay. Primers used to clone the different truncated version of RRP6L1are listed in Supplemental Table S4.

### Plant materials, growth conditions and physiological assays

The T-DNA insertion line of AHL10, *ahl10-1* (SALK_073165C) was previously described (Wong et al, 2019). T-DNA insertion lines of RRP6L1, *rrp6l1-1* (SALK_004432) and *rrp6l1-2* (SALK_027601) and the *ahl13-1* (SALK_014014) were obtained from the Arabidopsis Biological Resource Center and the homozygous plants confirmed using primers designed using the Signal web resource (http://signal.salk.edu/). To generate transgenic plants expressing *AHL10* under control of its native promoter, the genomic fragment containing the AHL10 gene body (5’ UTR, introns and exons up to the stop codon) and 2.1 kb of promoter upstream of the transcriptional start site were cloned into pENTR-D-TOPO and recombined into pGWB540 by Gateway recombination before transformation into *ahl10-1* or *ahl10- 1rrp6l1-2* (all in the Col-0 background) using *Agrobacterium tumefacians* strain GV3101 and the floral dip method. To express phosphomimic and phosphonull AHL10 under control of its native promoter, the entry clone in the pENTR-D-TOPO vector was used as template plasmid for PCR using site-directed mutagenic primers followed by Dpn1 digestion and cloning as previously described (Bhaskara et al., 2017). All constructs were verified by sequencing. Primers used for genotyping, cloning and site-directed mutagenesis are given in Supplemental Table S4.

Low ψ_w_ treatment on PEG-infused agar plates and soil drying experiments were performed as described previously (Verslues et al., 2006; Wong et al., 2019). For plate experiments to assay plant growth, 5-day-old seedlings were transferred from control media to the indicated treatment and growth measured 5 (control) or 10 (stress) days after transfer. For growth assay involving transgenic lines, three independent lines were tested in at least three independent experiments with two or three plates containing the test genotype and wild type used as technical repeats per experiment. Similar replication was performed for assays involving T-DNA mutants. Growth of the test genotype is reported relative to wild type growing in the same plate.

For soil drying experiments, a standard potting mix was combined with 25% Turface (Turface MVP, Profile Products LLC, USA). Four genotypes were planted in sectors (two plants per sector) of 8 cm x 8 cm x 10 cm (LxWxH) plastic pots to ensure that the different genotypes were exposed to the same extent of soil drying. Seeds placed into pots with fully watered soil mix, stratified and then transferred to a short-day chamber (8 h light period, 23 C, light intensity of 100-120 μmol m^-2^ sec^-1^) and Hyponex nutrient solution (1 g liter^-1^) supplied once per week along with regular watering to maintain full soil hydration. The position of individual pots within the chamber was rotated every two or three days. On day 19 after planting, pots were watered to saturation, allowed to drain and weighed. Water was then withheld for 12 days (leading to 50-60 percent reduction in pot weight), each pot re-watered to 75 percent of the initial pot weight and then allowed to dry another 8-10 day until pot weight again reached 50-60 percent of the starting weight. At the end of the experiment, representative rosettes were photographed and the rest weighed (fresh weight) and then dehydrated in an oven to obtain the dry weight. Growth of the test genotype is reported relative to wild type growing in the same pot. Each genotype was tested in at least three independent experiments with two or three pots containing the test genotypes and wild type used as technical repeats per experiment.

### In-vitro Pull-down assay

GST-tagged protein expression vector, pGEX-4T-1, was used to clone the RRP6L1 N-terminal 300 and C-terminal 337. AHL10 non-mutated wild type and phosphomimic of S313D and S314D was cloned into the pET300NT destination vector (N-terminal HIS tag). For GST-tagged RRP6L1 N- and C-terminal fusion proteins, the expression plasmids were transformed into BL21(DE3) pGro7 *E. coli* competent cells and induced by 0.5 mM isopropyl β-D-thiogalactoside (IPTG) in the presence of 0.5 g/L L-arabinose, and incubated for an additional 16-20 hours at 20°C. For His-tagged AHL10 (N.M., S313D and S314D), the expression plasmids were transformed into Rosetta 2(DE3) pLysS *E. coli* competent cells and induced by 0.3 mM IPTG, and incubated for an additional 22-24 hours at 16°C. The cells were harvested by centrifugation (6,000xg for 30 min at 4 [C) snap-frozen, and stored in - 80°C until use.

The bacterial pellets harboring GST-tagged RRP6L1 protein were resuspend in prechilled lysis buffer (20 mM HEPES-KOH at pH7.50, 200 mM NaCl, 0.1 mM MgCl_2_, 5% [v/v] glycerol, 5-mM 2-mercaptoethanol, 1x Complete, EDTA-free Protease Inhibitor Cocktail [Roche]), 1% [v/v] IGEPAL CA-630, 5 mM 2-mercaptoethanol), followed by sonication on ice using a 1/4-inch microtip controlled by the S-4000 ultrasonic processor (Misonix Sonicators). The cel lysates were clarified by high-speed centrifugation at 50,000g for 60 min at 4°C (Avanti J-26 XP centrifuge with JA-25.50 rotor; Beckman Coulter), followed by a 0.45-mm filtration (Millex-HP, 33 mm, polyethersulfone; Merck Millipore) before affinity chromatography. The pellets harboring His-tagged AHL10 protein were resuspend in prechilled lysis buffer (50 mM potassium phosphate at pH7.40, 200 mM NaCl, 10 mM Imidazole-Cl, 10% [v/v] glycerol, 5-mM 2-mercaptoethanol, 1x Complete, EDTA-free Protease Inhibitor Cocktail [Roche]), 5 mM 2-mercaptoethanol), followed by the same lysis and clarification procedures as RRP6L1.

For RRP6L1 N- and C-terminal fusion proteins (and free GST), GST affinity purification was performed using Glutathione Sepharose 4FF resin and proteins eluted using lysis buffer supplied additional freshly prepared 10 mM reduced glutathione-NaOH. His-tagged AHL10 proteins were purified using Clontech His60 Ni Superflow resin and eluted using 400 mM imidazole-Cl. The AHL10 proteins were rapidly diluted with equal volume of ion exchange buffer (10 mM potassium phosphate at pH7.40, 50 mM NaCl, 0.1 mM MgCl_2_, 10% [v/v] glycerol, 5-mM 2-mercaptoethanol) before being applied to a HiTrap Heparin HP (GE Healthcare), and eluted by a linear NaCl gradient up to 1.0 M. Purified proteins were dialyzed against pull-down buffer (25 mM HEPES [pH 7.5], 150 mM NaCl, 0.1 mM MgCl2, 10% [v/v] glycerol, 0.1% [v/v] IGEPAL CA-630, 5 mM 2-mercaptoethanol) and protein concentration measured using Bradford method (Bio-Rad Protein Assay Dye Reagent Concentrate; Bio-Rad Laboratories).

For pull down assays, 20 μl of glutathione magnetic beads (Pierce catalog no. 88821) were incubated with GST-tagged protein (30 μg free GST or 70 μg RRP6L1 N-300/C337) and 30 μg of AHL10 (10% of the reaction removed for input detection) for 120 min at 4 C with gentle rotation. Beads were then washed 5x with 1 ml of pull-down buffer and bound proteins eluted by adding 40 μl pull-down buffer and 10 μl of 5x SDS-PAGE Laemmli loading buffer to the washed beads. Proteins were then separated by SDS-PAGE and blotted onto PVDF membranes. Membranes were blocked using 5% nonfat milk and probed with anti-poly-His primary antibody (cat. no. H1029; Sigma-Aldrich) at 1:3,000 dilution or anti-GST primary antibody (cat. no. SAB4200692; Sigma-Aldrich) at 1:2,000 dilution, followed by secondary antibody (rabbit anti-mouse IgG H&L [HRP], AbCam ab6728) at 1:10,000 dilution). Immunoreactive bands were visualized by the enhanced chemiluminescence method (Pierce ECL Western Blotting Substrate, Thermo Fisher Scientific) and detected using Thermo Scientific CL-XPosure Film). For band quantification, images of the film (48- bit RGB tiff, 720dpi) were converted to 16-bit gray scale images. ImageJ was used to subtract background, invert the images and then measure intensity of equal sized regions of interest for each band. The ratio of pulldown band versus input band was calculated for each protein interaction and the data for unmutated AHL10 compared to the AHLS313D and AHL10S314D data. Three independent sets of pulldown reactions were conducted and used for band intensity quantification.

### Purification of His-tagged RRP6L1 C-terminal fragment and Generation of RRP6L1 antisera

The portion of the RRP6L1 cDNA encoding the C-terminal portion of the protein (amino acid 301-638) was cloned into pENTR-D-TOPO and LR reaction transferred to the pET300/NT-DEST vector to generate recombinant protein with an N-terminal fusion to 6×HIS tag. Recombination protein was produced in Rosetta 2(DE3) pLysS *E. coli* competent cells by addition of 1mM IPTG when cultures reached an optical density of 0.4 followed by incubation for 4 h at 37°C. Cells were harvested and resuspended in 50 mL lysis buffer (50 mM HEPES pH 7.4—adjusted pH with NaOH, 150 mM NaCl, 0.5% Tween-20, 1X protease inhibitor, 1 mM DTT, 15% glycerol) and lysed using a cell disruptor (Constant Systems).

Lysate was filtered and applied to a 1 mL Mini Profinity IMAC cartridge using BioLogic low-pressure chromatography system (Bio-Rad). An imidazole gradient was used to elute the protein in elution buffer (300 mM imidazole; 50 mM HEPES pH 7.4—pH adjusted with NaOH, 300 mM NaCl, 20% glycerol). Protein purify was checked by SDS-PAGE and commassie blue staining. For final purification, 2 mg purified protein was separated by SDS-PAGE and band corresponding to the expected molecular weight of the RRP6L1 301-638 fragment (40 kDa) excised and used for antisera generation in rabbit (performed by LTK Biolaboratories, Taiwan). Antibody titer and specificity was checked by immunoblot using protein extracts from the wild type and *rrp6l1* mutants probed with affinity purified RRP6L1 antisera (immunoblot procedures and optimized antisera dilution are described below).

### Ratiometric Bi-molecular Fluorescence Complementation (rBiFC) and subcellular localization

Ratiometric BiFC (rBiFC) was performed using the vector system described by (Grefen and Blatt, 2012) whereby a constitutively expressed RFP is present on the BiFc plasmid. AHL10 wild type, phosphomimic, or phosphonull cDNA clones (described in Wong et al., 2019) were inserted into pDONR221 P1-P4 (Invitrogen) and the RRP6L1 cDNA was amplified from total mRNA and cloned into pDONR221 P3-P2 (Invitrogen). The AHL27 and AHL29 cDNAs were amplified from plasmid clones generously provided by David Faverro (Riken) and cloned into pDONR221 P1-P4. Primer sequences used for cloning are given in Supplemental Table S4. These donor vectors were used for two site Gateway recombination into the destination vector pBiFCt2in1-NN and transformed into the Agrobacterium GV3101. Five-day-old seedlings of an Arabidopsis line with Dexamethasone (Dex) inducible AvrPto expression (Tsuda et al., 2012) were used for transient expression, followed by confocal microscopy (using a Zeiss LSM 510 META or Zeiss LSM 880), and quantification of YFP and RFP signal intensities performed as previously described (Wong et al., 2019). Subcellular localization of *AHL10_pro_:AHL10-EYFP*/*ahl10-1* and its phosphomimic and phosphonull variants were visualized using a Zeiss LSM 880 with airyscan.

### Collection of meristem-enriched tissue samples and immunoblotting

Seven-day-old seedlings were transferred to control and stress treatments (−0.7 MPa PEG-infused plates) for 4 days. Samples of the shoot meristem and surrounding tissue were collected using a 7mm diameter round metal punch to excise all tissue surrounding the meristem regions. Root tip samples were collected by excising the apical 3-5 mm of the root tip with a razor blade. Meristem-enriched samples or whole seedlings were ground in liquid nitrogen and homogenized in cold protein extraction buffer (50 mM Tris HCl pH 7.5, 10% glycerol, 1 mM DTT, 1% IGEPAL, 1× Roche EDTA-free protease inhibitor and 1× Roche phosphatase inhibitor). The homogenate was centrifuged and the protein concentration of the supernatant was measured using a Pierce 660 nm Protein Assay Reagent (Thermo Scientific). Thirty-five microgram of protein was loaded in each lane. Two SDS-PAGE gels were run with the same aliquots of all the tissues samples, one was used for anti-GFP (Sigma-Aldrich Cat# 1181446001) and the other used for blotting with anti-RRP6L1 polyclonal antisera described above (antibody dilutions and other blotting conditions are described below). The anti-RRP6L1 blots were stripped using mild stripping buffer (1.5 g glycine, 0.1g SDS, 1 mL Tween-20 in 100 mL solution, pH 2.2) and re-probed with anti-HSC70 (Enzo # ADI-SPA- 818-D) at 1:5000 dilution, as a loading control.

### Isolation of total nuclear, nucleoplasmic and chromatin-associated protein

Seven-day-old seedlings were transferred to -0.7MPa PEG-infused plates for 4 days. Seedlings were harvested and cross-linked with 1% formaldehyde solution in 1× PBS by vacuum infiltration (3 times, 10 minutes total infiltration time with total exposure time to formaldehyde not exceeding 15 minute). Cross-linking was stopped by adding 0.125 M glycine followed by vacuum infiltration (2 times, 90 seconds each time). The samples were rinsed twice with cold PBS, dried with a paper tower and 600 mg of tissue frozen in liquid nitrogen. The same cross-linking procedure was used for assays of chromatin associated protein as well as ChIP-quantitative PCR assays (described below).

Isolation of total nuclear, nucleoplasmic and chromatin-bound associated was conducted following the procedure of Wu et al., (2016) with some modifications. Samples were ground into powder using a mortar and pestle and suspended in 5 ml of Honda Buffer (20 mM HEPES-KOH pH 7.4, 0.44 M sucrose, 1.25% ficoll, 2.5% Dextran T40, 10 mM MgCl2, 0.1% Triton X-100, 5 mM DTT, 0.5 mM PMSF, 1× protease inhibitors [Roche]), filtered through two layers of Miracloth and centrifuged at 2000 x g for 15 min at 4 °C. The supernatant was removed, and the nuclear pellets were washed twice with 1ml of Honda buffer, resuspended in 600 μl nuclei lysis buffer (10 mM Tris-HCl pH 7.5, 2 mM EDTA, 1× proteinase inhibitor, 1× phosphatase inhibitor and 1% SDS). 600 μl aliquots of lysed nuclei solution were aliquoted into two low binding microcentrifuge tubes. One tube was kept on ice (this is the total nuclear extract), and the other tube vortexed for 2 minutes followed by centrifugation at 14,000g for 10 mins at 4°C and collection of the supernatant (this is the soluble fraction of nucleoplasm protein). The remaining pellet was re-suspended in 300 µl of lysis buffer containing 1% SDS and sonicated using a bioruptor (Diagenode) at 4°C (15 sec ON/OFF for 5-min and repeated for 5 cycles). The sonicated fragment was centrifuged at 14,000g for 10 mins at 4°C and the supernatant collected (this is the chromatin-associated protein). The pellet was resuspended and collected (pellet after sonication fraction). From the collected fractions of total nuclear extract, soluble nucleoplasm, chromatin-associated and pellet after sonication (each in 300 μl total volume), 50 μl was mixed with 6× loading dye, heated at 95°C for 10 mins, loaded on 10% SDS-PAGE, and transferred to PVDF membrane followed by blocking in 5% w/v non-fat milk powder in 1% TBST for overnight. The anti-GFP antisera (Sigma-Aldrich Cat# 1181446001) was used at 1:2000 dilution followed by 1:10000 secondary anti-mouse HRP. The membrane was stripped using mild stripping buffer (1.5 g glycine, 0.1g SDS, 1 ml Tween-20 in 100 ml solution, pH to 2.2) and re-probed with anti-RRP6L1 antibody (1:8000 dilution) followed by 1:10000 anti-rabbit HRP. Separate 50 µl (same aliquot from the total 300 µl) were separated on 13% SDS-PAGE gel for immunoblot using anti-Histone 3 (Sigma #06-755) at 1:3000 dilution with 1:10000 anti-rabbit secondary antibody. Quantification of band intensity for the 74 kd RRP6L1-specific band as well as AHL10-YFP band was performed as described above for the GST pull down experiments.

### ChIP-quantitative PCR assays

ChIP-quantitative PCR (qPCR) assays were performed as previously described (Yamaguchi et al., 2014). Plant samples were collected, harvested and cross-linked as described above for the chromatin-associated protein assays. 300 mg tissue from whole seedlings was used with 30 µl of GFP magnetic trap beads (Chromotek). The plants were ground to powder in liquid nitrogen and lysed in 2.5 ml cold nuclei extraction buffer. The lysate was filtered through 2-layers of miracloth (Millipore, Cat. 475855) set up in a 50[ml centrifuge tube, centrifuged at low speed to collect the pellet (nuclei) and the pellet resuspended with nuclei lysis buffer. Sonication was performed in an ice bath using a Bioruptor (Diagenode) set to high power and operated using 15sec on/off cycles for 5-min and repeated for 3 cycles. The procedures for nuclei isolation, shearing of chromatin, immunoprecipitation, washing, reverse crosslinking, elution to get the purified DNA using QIAquick spin columns (Qiagen, Cat: 28106) and ChIP-qPCR DNA reaction composition were as described in Yamaguchi et al. (2014). A 5-fold dilution series of input was amplified for each set of primers and the data used to calculate the percent input. Two or three independent experiments were performed for each ChIP target site analyzed, as described in the figure legends. Primers used for ChIP-qPCR assays are shown Supplemental Dataset S4.

### RNA Sequencing and Quantitative Reverse Transcriptase-PCR analysis of gene expression

The RNA sequencing of *rrp6l1-2* was conducted in the same set of experiments as previously reported data of wild type and *ahl10-1* and followed the previously described procedures for sample processing, and data analysis (Wong et al. 2019). Seven-day-old seedlings were transferred to fresh control plates or PEG-infused agar plates (−0.7 MPa) and whole seedlings collected four days after transfer (11 days old). Significant differences in gene expression were determined using DEseq2. Heat maps with hierarchical clustering (Euclidian distance clustering) were prepared using Morpheus (Broad Institute, <u>Morpheus</u> (broadinstitute.org)).

For QPCR, the sample collection procedure was that same as described above for RNAseq analysis. Total RNA was extracted from control or -0.7 MPa stress-treated seedlings using RNeasy Plant Mini Kit (Qiagen). 1 μg RNA was reverse transcribed using SuperScript III (Invitrogen). Real-time quantitative PCR assays were prepared using KAPA SYBR FAST qPCR kit (Kapa Biosystems) and run on an Applied Biosystems QuantStudio 12K QPCR instrument. The comparative ΔΔC_T_ method was used to determine relative gene expression using *EFL1α* as a reference gene (Nicot et al., 2005). Note that for some genes (*STM, WES1*, *DFL1, MYB117* and *PRR*) the QPCR data *ahl10-1* was previously reported in Wong et al. (2019) and is shown again here for comparison to *rrp6l1-2*.

### Statistical Analysis

Graph preparation and statistical analyses were performed using GraphPad Prism 9. Data were analyzed by ANOVA, T-test or one sample T-test as indicted in figure legends with corrected P ≥ 0.05 used as a cutoff for significant differences.

## Results

### AHL10-interacting proteins, including RRP6L1, identified by yeast two-hybrid screening

To identify AHL10 interacting proteins, we used full length AHL10 as bait to screen a yeast two-hybrid library prepared using cDNA isolated from low ψ_w_-treated seedlings (the library has been described in Chong et al., 2019). The screening identified several clones of AHL1, AHL3, AHL10 and AHL13 that showed strong interaction with AHL10 (Supplemental Fig. S1A). This indicated that these four Clade B AHLs can form complexes with each other and is consistent with the results of Jiang et al. (2017) where TAP tagging and immunoprecipitation of ADM also found these four AHLs. Among other nuclear-localized proteins identified in the screen, AHL10 strongly interacted with RRP6L1 and the cell-cycle regulator CDC20.2 and had weaker putative interaction with RNA-binding (RBP45A), DNA-binding (WRKY1, Histone 3.1) and chromatin associated (SGO2) proteins. We focused further attention on RRP6L1 because of its strong interaction in the yeast-two hybrid assays (Fig. 1A). The interacting clone detected in the yeast two-hybrid screen contained the C-terminal 76 amino acids of RRP6L1. Retransformation assays confirmed AHL10 interaction of this clone as well as a longer clone containing the C-terminal 192 amino acids. However, neither the C-terminal 338 amino acids nor full length RRP6L1gave a positive interaction signal with AHL10 (Fig 1A). It is possible that the longer RRP6L1 clones interfered with reporter activation in the yeast assays.

**Figure 1:**
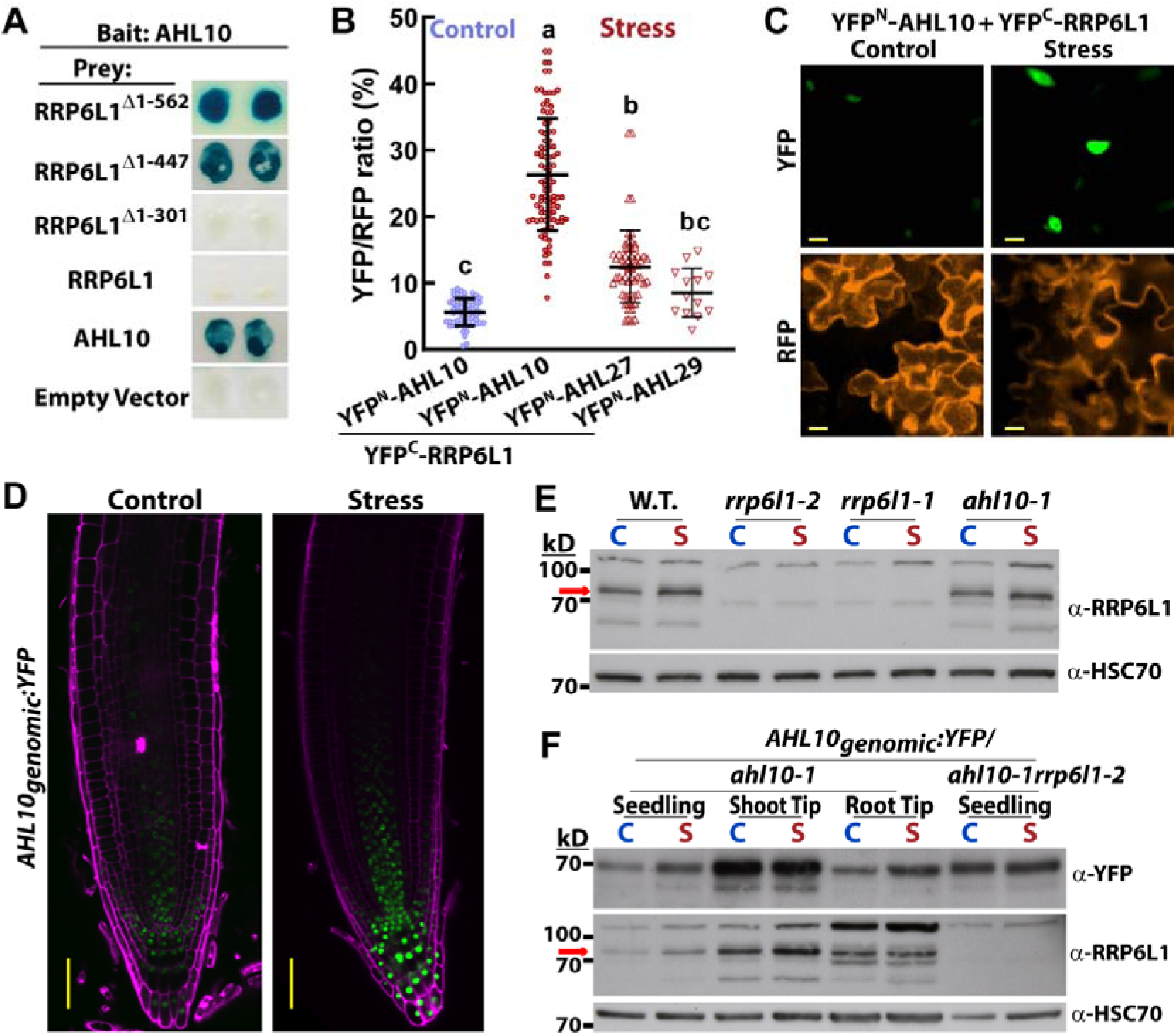
AHL10 interacts with RRP6l1 and both proteins accumulate in root and shoot meristem regions. A. β-galactosidase staining colony lift assays show the interaction of AHL10 with clones encoding the C-terminal portion of RRP6L1. RRP6L1^Δ1-562^ is the clone found in the original yeast two hybrid library screening. Other truncations and full length *RRP6L1* clones were used to confirm the interaction. AHL10 self-interaction is shown for comparison as another strong interaction detected in the yeast two hybrid screen. Empty prey vector is a negative control. Additional proteins found in the yeast two hybrid screen are listed in Supplemental Fig. S1A. B. Quantification of interaction intensity for rBIFC assays conducted using unstressed Arabidopsis seedlings or seedlings exposed to low ψ_w_ stress (−0.7 MPa for 48 h). Experiments were conducted using transient expression in Arabidopsis seedlings with constructs driven by the *35S* promoter (see methods). Individual data points represent the YFP/RFP fluorescence intensity ratio of individual cells. Data are combined from three independent experiments (n = 13 - 97). Different letters indicate statistically significant differences (ANOVA, corrected P ≤ 0.05). C. Representative images of rBiFC data presented in B. Scale bars indicate 10 μm. D. Localization of AHL10-YFP expressed under control of its native promoter (genomic clone of the *AHL10* gene body consisting of 5’ UTR, introns and exons up to the stop codon, and 2.1 kb promoter region; with C-terminal YFP fusion). Images show the primary root tip of 9-day-old seedlings growing control media or four days after transfer to low ψ_w_ (−0.7 MPa). Scale bars indicate 50 μm. E. Validation of polyclonal antisera raised against the C-terminal region of RRP6L1. Red arrow marks major band corresponding to the molecular weight of RRP6L1 (74 kD) which is absent in both *rrp6l1* mutants. 35 μg of protein was loaded in each lane. A non-specific band was detected at approximately 110 kD. The blot was stripped and re-probed with anti-HSC70 as a loading control. The experiment was repeated with essentially identical results. C = Control; S = Stress (−0.7 MPa for four days). The blot was stripped and re-probed with anti-HSC70 as a loading control. F. Detection of RRP6L1 and AHL10-YFP in whole seedling, shoot meristem-enriched (shoot tip), and root meristem-enriched (root tip) samples. 35 μg of protein was loaded in each lane. The RRP6L1 blot was stripped and re-probed with anti-HSC70 as a loading control. The experiment was repeated three times with consistent results each time.

### AHL10-RRP6L1 interaction

Ratiometric Bi-molecular fluorescence complementation (rBiFC) assays found that AHL10-RRP6L1 generated a strong fluorescence signal in seedlings exposed to low ψ_w_ stress but not in unstressed seedlings (Fig. 1B and C). As a control for the rBiFC interaction assays, we also tested AHL10 interaction with AHL27 and AHL29. In the low ψ_w_ treatment, AHL27 and AHL29 had lower rBiFC signal compared to the AHL10-RRP6L1 interaction (Fig. 1B). This lower rBiFC signal may represent non-specific background, perhaps due to high level of transient expression of the proteins, or may represent a limited interaction of RRP6L1 with other AHLs. We also found that AHL10 had a relatively strong rBiFC signal with WRKY1 in both unstressed and low ψ_w_-treated plants (Supplemental Fig. S1B), indicating that AHL10 expression was not the cause of the lower AHL10-RRP6L1 rBiFC signal in unstressed plants. Together these data demonstrated that RRP6L1 interacted with AHL10 in the nucleus and that this interaction may be stimulated by low ψ_w_ stress.

AHL10-RRP6L1 interaction was also detected using *in vitro* pull-down assays. RRP6L1 was divided into the N-terminal 300 amino acids (N300) and C-terminal 337 amino acids (C337) as full length RRP6L1 could not be expressed in *E. coli*, consistent with previous report (Zhang et al., 2014). The N300 fragment of RRP6L1 contains the 3’-5’ exonuclease domain while the C337 fragment contains a Helicase and RNase D C-terminal (HRDC) nucleic acid-binding domain. Interestingly, both halves of RRP6L1 interacted with AHL10; however, the pull-down band intensity was significantly greater for the C337 fragment of RRP6L1 than for N300 (Supplemental Fig. S2A and C). Phosphomimic mutations at the S313 and S314 phosphorylation sites previously found to be critical for AHL10 function (Wong et al., 2019) increased the pull-down band intensity of the N300 fragment but had no significant effect on interaction of C337 (Supplemental Fig. S2B and C). These data confirmed AHL10-RRP6L1 interaction and indicated that AHL10 phosphorylation at S313 or S314 is unlikely to block interaction with RRP6L1; however, we cannot rule out that phosphorylation affects the affinity of AHL10-RRP6L1 interaction or affects how AHL10 interacts with the different domains of RRP6L1.

### AHL10 and RRP6L1 accumulate in meristematic tissue

We generated transgenic lines where a genomic fragment containing the AHL10 promoter and gene body (5’ UTR, exons, and introns) up to the stop codon was fused with YFP (*AHL10_genomic_:YFP*) and used to complement *ahl10-1* (Supplemental Fig. S3). Imaging of these transgenic lines found that AHL10-YFP accumulated in the root quiescent center, developing stele and columella cells under both control and low ψ_w_ stress conditions (Fig. 1D). AHL10-YFP also accumulated to high levels in the shoot meristem region as well as in the meristem region of lateral roots and sites of lateral root emergence (Supplemental Fig. S4). The same AHL10 native promoter construct was produced for phosphonull AHL10 (AHL10^S313A^, AHL10^S314A^) and phosphomimic AHL10 (AHL10^S313D^, AHL10^S314D^) and transformed into *ahl10-1*. Consistent with previous results AHL10^S313D^ and AHL10^S314D^ complemented *ahl10-1* while AHL10^S314A^ did not complement. AHL10^S313A^ was hyper- functional and suppressed growth below the wild type level during low ψ_w_ stress (Supplemental Fig. S3). We hypothesize that preventing phosphorylation at S313 caused S314 to become hyperphosphorylated (Wong et al 2019). Hyperphosphorylation of S314 may have larger growth suppression effect than the S314 phosphomimic construct as aspartate substation, while useful for most functional analyses, is not a perfect mimic of phosphorylation (it introduces a single negative charge while phosphorylation introduces a -2 charge). AHL10^S314A^ was diffusely localized in the nucleoplasm and lacked the nuclear foci seen for wild type AHL10, AHL10^S313A^ and phosphomimic AHL10 (Supplemental Fig. S5). Note that the nucleoplasm localization of AHL10-YFP matches previously reported localization of RRP6L1 (Lange et al., 2008; Zhang et al., 2014).

To study RRP6L1 protein accumulation, and compare it to that of AHL10, we generated antisera recognizing the N-terminal domain of RRP6L1. Immunoblotting of wild type and two previously described RRP6L1 knockout mutants, *rrp6l1-1* (Lange et al., 2008; Zhang et al., 2014; Ye et al., 2016; Wendte et al., 2017) and *rrp6l1-2* (Hsu et al., 2014), found that the antisera recognized a 74 kD band which corresponded to the expected molecular weight of RRP6L1 and which was absent in both of the *rrp6l1* alleles (Fig. 1E). It was also consistently observed that RRP6L1 protein level was moderately increased by low ψ_w_ stress. Samples of shoot and root meristem (and surrounding tissue) had substantially higher levels of RRP6L1 and AHL10-YFP than whole seedling samples (Fig. 1F). These data indicated that AHL10 and RRP6L1 proteins accumulate in the same parts of the plant, consistent with their physical interaction. Immunoblots also showed that RRP6L1 protein level did not substantially differ between wild type and *ahl10-1.* Likewise, AHL10 protein level was not substantially affected by *rrp6l1-2* (Fig. 1E and F). Thus, the purpose of the AHL10-RRP6L1 interaction was not related to one protein affecting the abundance of the other protein.

### RRP6L1 and AHL10, but not AHL13, restrict growth at low **ψ**_w_

We subjected both *rrp6l1* alleles to a moderate severity low ψ_w_ (−0.7 MPa) that decreases, but does not completely inhibit, growth of wild type (Wong et al., 2019). Both *rrp6l1* mutants maintained a higher level of growth than wild type during this moderate severity low ψ_w_ stress while having minimal or no effect on growth in the unstressed control, similar to *ahl10-1* (Fig. 2A, Supplemental Fig. 6A, wild type data used for normalization can be found in Supplemental Fig. S6B). An *ahl10-1rrp6l1-2* double mutant had similar increased growth maintenance as either single mutant. *rrp6l1-2* also had enhanced growth maintenance during controlled soil drying assays where plants were exposed to an extended period of partial soil drying that slowed rosette growth but did not lead to wilting (Fig 2B, C). Note that the *ahl10-1* data from these soil drying experiments has been previously reported (Wong et al., 2019) and is shown again here to allow better comparison between all genotypes.

**Figure 2:**
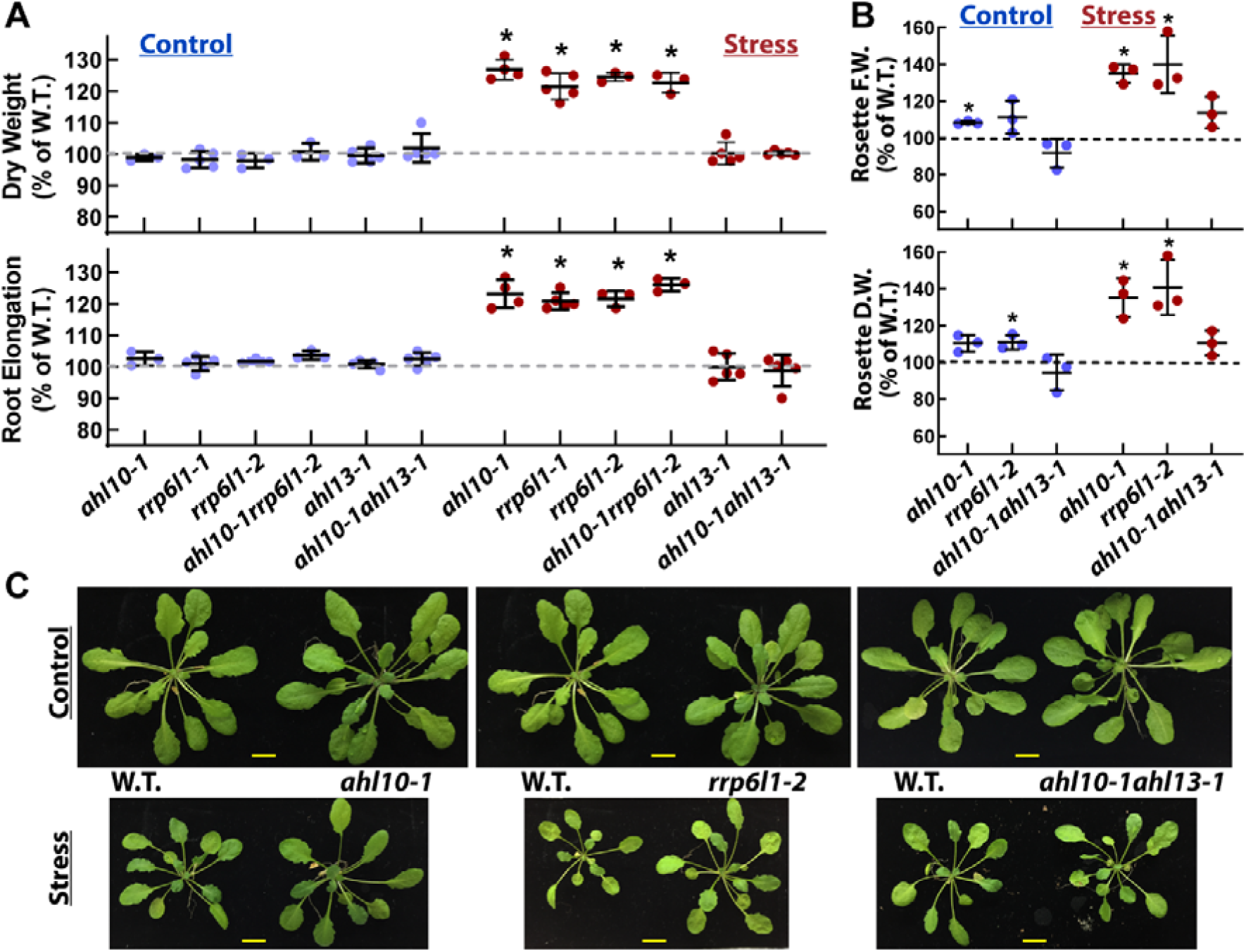
*RRP6L1* is a negative regulator of growth during moderate severity low ψ_w_ stress and *ahl10-1* effect on growth depends upon *AHL13*. A. Seedling dry weight and primary root elongation in control and stress treatments. For control, five-day-old seedlings were transferred to fresh control agar plates with root length increase and total seedling dry weight measured five days after transfer. For stress treatment, five-day-old seedlings were transferred from control media to PEG-infused agar plates (−0.7 MPa) and dry weight and root elongation measured 10 days after transfer. Data are expressed relative to wild type grown in the same plate as the test genotype. Pictures of representative seedlings in the stress treatment are shown in Supplemental Fig. S6A. Wild type dry weight and root elongation data used for normalization are shown in Supplemental Fig. S6B. Data are means ± S.D. for 3-5 replicate experiments with 2-3 plates of seedlings measured in each experiment. Asterisks (*) indicate significant difference from 100% (wild type level) by one sample T-test. B. Rosette fresh and dry weights for soil-grown plants kept under well-watered conditions or exposed to partial soil drying. Data are expressed relative to wild type plants grown together in the same pot. Data are means ± S.D. for 3-5 replicate experiments (2-3 biological replicates per experiment). Asterisks (*) indicate significant difference from 100% (wild type level) by on sample T-test. C. Pictures of representative wild type and *rrp6l1-2* plants from the experiments presented in B. Scale bars indicate 1 cm.

AHL13 is closely related to AHL10, including conservation of the C-terminal phosphorylation site (Wong et al., 2019; Rayapuram et al., 2021), and likely forms heteromeric complexes with AHL10 (Supplemental Fig. S1). However, *ahl13-1* did not differ from wild type in growth maintenance at low ψ_w_ (Fig 2A). Interestingly though, loss of AHL13 blocked the increased growth of *ahl10-1* as *ahl10-1ahl13-1* grew similarly to wild type in both PEG-plate (Fig. 2A) and soil drying assays (Fig 2B). Transformation of *ahl10- 1ahl13-1* with *35S:AHL13* restored the *ahl10-1* phenotype (Supplemental Fig. 6C), validating that AHL13 is required for the increased growth maintenance of *ahl10-1*. These data indicate that AHL10 and AHL13 do not have redundant function. Rather AHL13 acts genetically downstream of AHL10, consistent with the formation AHL10-AHL13 heterocomplexes.

### AHL10 S314 phosphorylation is required for RRP6L1 chromatin association

AHLs and RRP6L1 are both hypothesized to require additional interacting factors to correctly target and associate with specific genomic sites or transcripts (Zhao et al., 2013; Wendte et al., 2017; Favero et al., 2020). Thus, we used chromatin precipitation to test whether AHL10 or RRP6L1 affected one another’s partitioning to the nucleoplasm or chromatin-associated fractions of nuclear extracts and whether this partitioning was affected by low ψ_w_ stress. In these assays, centrifugation of nuclear extracts was used to separate soluble nuclear proteins (Soluble Fraction, labeled as “S.F.” in Fig. 3) from chromatin-associated proteins (labeled as “C.A.” in Fig. 3). Sonication then allowed the chromatin-associated proteins to be solubilized and separated from the remaining insoluble material (pellet after sonication, labeled as “P.S.” in Fig. 3).

**Figure 3:**
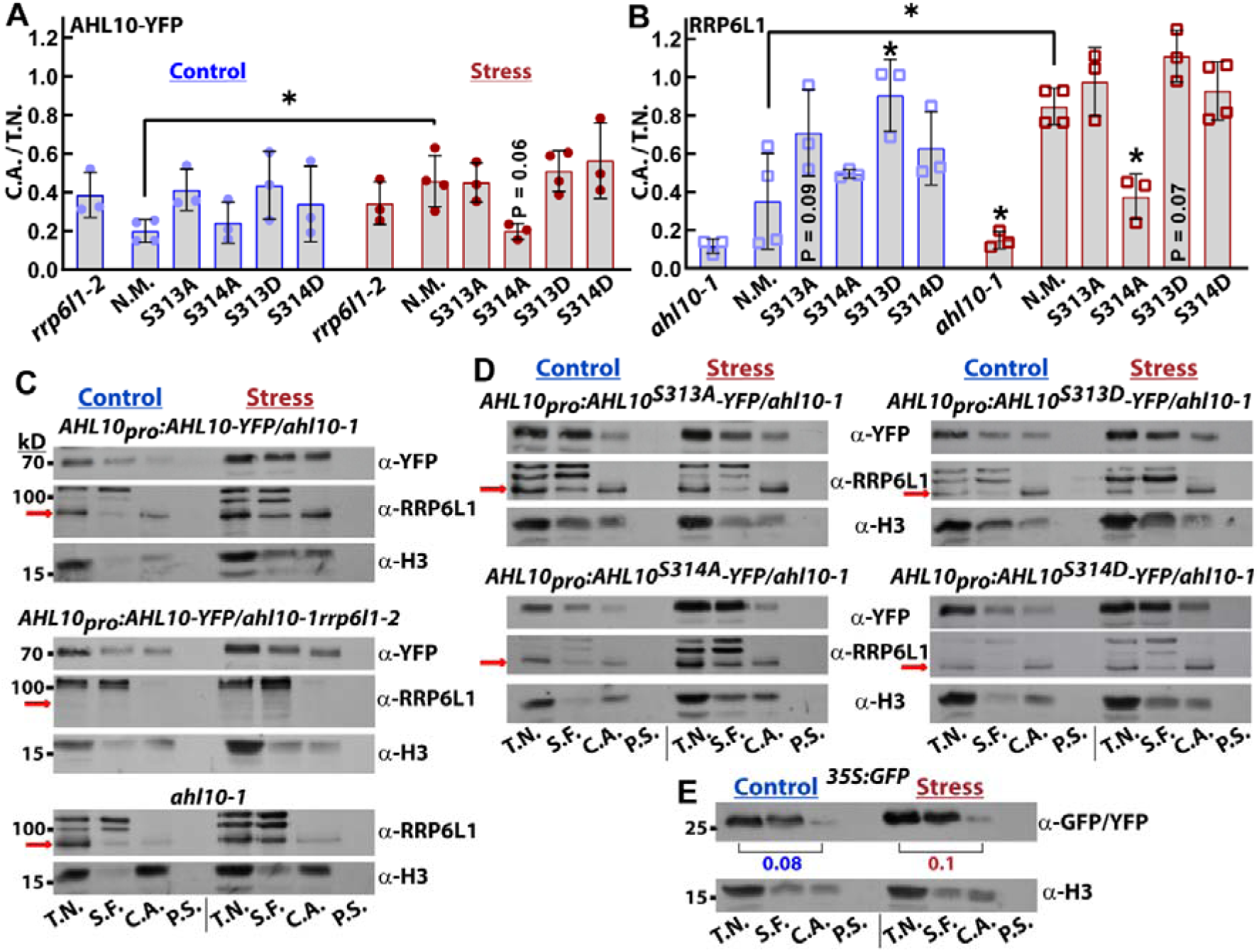
RRP6L1 chromatin association depends upon AHL10 S314 phosphorylation. A. Portion of AHL10-YFP associated with chromatin, measured as the ratio of chromatin associated (C.A.) versus total nuclear (T.N.) immunoblot band intensities for *rrp6l1-2* or *ahl10-1* complemented with non-mutated AHL10 (N.M), phosphonull AHL10 (S313A, S314A) or phosphomimic AHL10 (S313D, S314D). Representative blots used for quantitation can be seen in C and D. Samples were collected for 11-day-old seedlings kept under unstressed control conditions or transferred to low ψ_w_ stress treatment (−0.7 MPa for four days). Data are means ± S.D. for three or four independent experiments (one replicate per experiment). Asterisks or indicated P value at the bars in the graph indicate the results of ANOVA comparing all genotypes within the stress or control treatments to the wild type AHL10 (N.M.) data for that treatment. The wild type AHL10 (N.M.) data were further compared between control and stress treatment using T-test (Asterisk indicates significant difference, P ≤ 0.05). B. Portion of RRP6L1 associated with chromatin measured as the ratio of chromatin associated (C.A.) versus total nuclear (T.N.) immunoblot band intensities for *ahl10-1* or *ahl10-1* complemented with non-mutated AHL10 (N.M), phosphonull AHL10 (S313A, S314A) or phosphomimic AHL10 (S313D, S314D). Representative blots used for quantitation can be seen in C and D. Sampling conditions and statistical analysis are as described in A. Asterisks above bars indicate a significant difference compared to AHL10 N.M. in that treatment (P ≤ 0.05). C. Representative blots used for band intensity quantitation shown in A and B. The AHL10 genomic fragment with 5’ UTR and 2.1 kb promoter region was fused to YFP and expressed in the *ahl10-1* or *ahl10-1rrp6l1-2* genetic backgrounds to detect AHL10 chromatin association in the presence of absence of RRP6L1. The *ahl10-1* mutant was used to detect RRP6L1 chromatin association in the absence of AHL10. Red arrow indicates the RRP6L1 specific band which matches the predicted RRP6L1 molecular weight (74 kD) and was used for quantification. T.N. = Total Nuclear; S.F. = Soluble Fraction of nuclear proteins; C.A. = Chromatin Associated; P.S. = Pellet after Sonication (to check for remaining insoluble protein). Detection of Histone3 (H3) was performed as a control for the purity of the S.F. and C.A fractions. However, note that we consistently observed that stress treatment caused an increase in the amount of H3 in the soluble fraction which was not related to any technical aspects of the chromatin precipitation procedure. D. Representative blots of lines expressing phosphomimic or phosphonull AHL10 used for band intensity quantification shown in A and B. E. Nuclear fractionation and chromatin association of a transgenic line expressing GFP under control of the constitutive *35S* promoter used to check the background level of a non-chromatin associated protein in the chromatin pellet. Numbers under the GFP blot indicate the fraction of chromatin associated GFP based on the ratio of C.A and T.N. band intensities. Data are representative of two biologically independent experiments.

RRP6L1 presence in the chromatin-associated fraction was strongly increased by low ψ_w_ and was dependent upon AHL10 S314 phosphorylation. RRP6L1 chromatin association was reduced to background level in *ahl10-1* (compare Fig. 2B and C to Fig. 2E where a line expressing GFP alone was used as a negative control). In the low ψ_w_ treatment, RRP6L1 presence in the chromatin-associated fraction was also strongly reduced by the AHL10^S314A^ phosphonull mutation (Fig 2B and D). This reduced RRP6L1 chromatin association in plants expressing AHL10^S314A^ correlates with the altered sub-nuclear localization of AHL10^S314A^ (Supplemental Fig. S5) and with its inability to complement *ahl10-1* (Supplemental Fig. S3). Interestingly, the opposite effect was observed for AHL10^S313A^ where the portion of RRP6L1 in the chromatin-associated fraction was high and similar to the high level seen in the AHL10 phosphomimic mutants (AHL10^S313D^ and AHL10^S314D^). The high level of chromatin-associated RRP6L1 in plants expressing AHL10^S313A^ correlates with the strong growth suppression activity of AHL10^S313A^ (Supplemental Fig. S3; Wong et al., 2019) and is consistent with the idea that blocking AHL10 S313 phosphorylation directs more phosphorylation to the critical S314 site (Wong et al., 2019). These effects of AHL10 on RRP6L1 chromatin association were more apparent in the low ψ_w_ stress treatment, consistent with the low ψ_w_-induced increase of AHL10-RRP6L1 association seen in rBiFC assays (Fig. 1B).

Conversely, the proportion of wild type AHL10 (labeled as N.M., for Non-Mutated) in the chromatin-associated fraction was not affected by *rrp6l1-2* but was significantly increased by low ψ_w_ stress (Fig. 3A and C). We also found that the AHL10^S313A^, AHL10^S313D^, and AHL10^S314D^ phosphomimic and null mutations did not affect AHL10 presence in the chromatin-associated fraction (Fig. 3A and D), consistent with phosphorylation of the C-terminal domain having no direct effect on DNA binding activity of the N-terminal AT-hook domain. AHL10^S314A^ may have been somewhat less present in the chromatin-associated fraction at low ψ_w_, however, the effect was marginally non-significant (P = 0.06) and thus should be interpreted with caution.

The remaining pellet after chromatin-associated proteins were released by sonication did not contain detectable AHL10 or RRP6L1, suggesting that neither protein was part of other, non-chromatin-associated, insoluble protein aggregates in the nucleus. Histone 3 (H3) was used as a control chromatin-associated protein (Fig. 3C). Note that our assays indicated that low ψ_w_ stress led to an increased level of H3 in the soluble nuclear fraction but had little effect on the chromatin associated fraction. This indicated that the increased chromatin association of AHL10 and RRP6L1 in the stress treatment was a specific effect and not a general effect of the low ψ_w_ treatment on all nuclear proteins.

### RRP6L1 and AHL10 affect the expression of an overlapping set of low **ψ**_w_-responsive genes

We conducted transcriptome analysis of *rrp6l1-2* under unstressed conditions and after four-day acclimation to moderate severity low ψ_w_ stress (−0.7 MPa). This relatively long term low ψ_w_ treatment was used to find genes that undergo sustained induction or repression and thus are more likely to be associated with the differences in growth between *rrp6l1* versus wild type at low ψ_w_. The *rrp6l1-2* transcriptome data were collected in the same set of experiments as *ahl10-1* and wild type data previously reported (Wong et al., 2019). In comparison of unstressed wild type to unstressed *rrp6l1-2*, we found 147 genes having increased expression and 147 genes having decreased expression, including *RRP6L1* itself (log2 fold-change ≥ 0.5, adjusted P ≤ 0.05; Supplemental Table S1). Genes up- or down-regulated in *rrp6l1-2* had the same direction of change in *ahl10-1* in nearly all cases (Supplemental Fig. S7A), albeit that the *ahl10-1* effect was weaker in many cases. This was consistent with AHL10 and RRP6L1 acting together in gene regulation. Many of the genes with altered expression in unstressed *rrp6l1-2* were stress responsive in wild type (Supplemental Fig. S7A). Particularly, genes with increased expression in unstressed *rrp6l1- 2* were enriched for GO terms related to abiotic stress or environmental stimuli (Supplemental Table S2), suggesting that RRP6L1 has a role in preventing aberrant expression of these genes in unstressed conditions.

In the low ψ_w_ stress treatment, relatively fewer genes were differentially expressed in *rrp6l1-2* (11 genes had increased expression, 11 genes had decreased expression including *RRP6L1*; log2 fold-change ≥ 0.5 versus wild type at low ψ_w_, adjusted P ≤ 0.05; Supplemental Table S3). There was again substantial concordance between gene expression changes seen in *rrp6l1-2* and *ahl10-1* (Fig. 4A). This concordance could be seen even more clearly if we considered a slightly relaxed cutoff for differential expression (log2 fold-change ≥ 0.3, adjusted P ≤ 0.05) as shown in Supplemental Fig. S7B. The closest concordance between *rrp6l1-2* and *ahl10-1* was for genes that had decreased expression in both mutants (Fig. 4A; Supplemental Fig. S7B). We validated the decreased expression of a number of these genes in low ψ_w_-treated *rrp6l1-2* and *ahl10-1* by QPCR (Fig. 4B). Interestingly, given the proposed functions of *rrp6l1-2* (Wendte et al., 2017), *RNA-Dependent RNA Polymerase 3* (*RdR3*) and *RdR4* had decreased expression in both *rrp6l1-2* and *ahl10-1*. These two RdRs are of unclear physiological function (Willmann et al., 2011). The large number of genes with relatively small change in expression in *rrp6l1-2* and *ahl10-1* (Supplemental Fig. S7B) was not unexpected given the tissue specific accumulation of both proteins (Fig. 2) such that localized changes in gene expression may have been diluted in the whole seedling samples used for RNAseq.

**Figure 4:**
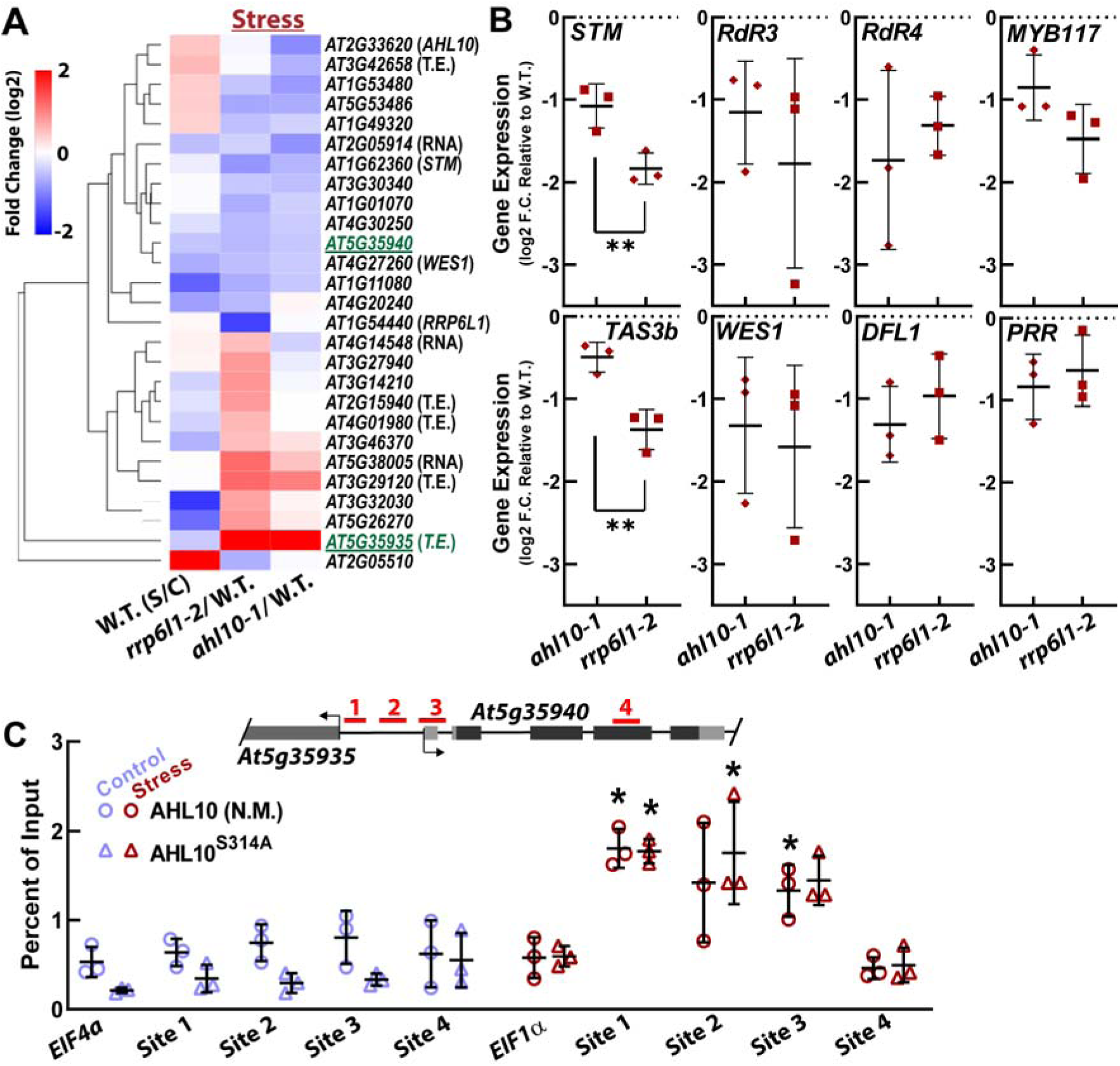
*rrp6l1-2* and *ahl10-1* have concordant effects on gene expression and low ψ_w_ stress can increase association of AHL10 with specific AT-rich regions. A. Heat map and hierarchical clustering of genes having significantly increased or decreased expression (log2 fold change ≥ 0.5, corrected P-value ≤ 0.05) in stress- treated (transfer to -0.7 MPa for four days) seedlings. Effect of stress treatment on these genes in wild type is also shown for comparison. Underlined genes were used as targets of ChIP assay shown in Fig. 4C. Similar heat map of genes with altered expression in unstressed *rrp6l1-2* or *ahl10-1* is shown in supplemental Fig. S7A. Heat map of genes affected by stress in *rrp6l1-2* or *ahl10-1* at slightly less stringent cutoff (log2 fold change ≥ 0.3, corrected P-value ≤ 0.05) is shown in Supplemental Fig. S7B. Genes labeled as “RNA” are non-coding RNAs. T.E. = Transposable Element. B. QPCR validation of gene expression differences of *rrp6l1-2* and *ahl10-1* compared to wild type in the stress treatment. Note that some of the genes selected for validation did not meet the cutoff for inclusion in the heat map shown in Fig. 4A but are included in the heat map shown in Supplemental Fig S7B. Data are means ± S.D. for data from 3 independent biological experiments (one sample per experiment, three technical PCR replicates per sample). Data are shown as log2 fold change relative to wild type. Asterisks indicate significant difference between *ahl10-1* and *rrp6l1-2* by T-test (**, P ≤ 0.01). Note that for some genes (*STM, WES1, DFL1, MYB117* and *PRR*) the QPCR data *ahl10-1* was previously reported in Wong et al. (2019) and is shown again here to facilitate comparison to *rrp6l1-2*. C. ChIP assays of AT-rich sites lying in the region between *At5g35935* and *At5g35940* (*At5g35940* promoter region) as well as an off-site in exon of *At5g35940* and a distal offsite (*EIF4a* promoter). Samples were collected for unstressed and low ψ_w_ -treated (−0.7 MPa for four days) seedlings. Data are means ± S.D. for data from 3 independent biological experiments (one sample per experiment, three technical PCR replicates per sample). Asterisks indicate significant differences compared to the *EIF1α* offsite in the stress treatment (ANOVA, P ≤ 0.05). No significant differences were found in the unstressed control.

AHL10 has been previously associated with AT-rich transposons (Jiang et al., 2017). Our transcriptome analysis of *ahl10-1* found altered RNA levels of several transposons (Wong et al., 2019) and many of these had similarly altered expression in *rrp6l1-2*. For example, the copia-like retrotransposon *At5g35935* and hAT-like transposase family gene *At3g29120* were the two most highly upregulated genes in the low ψ_w_ treatment for both *ahl10-1* and *rrp6l1-2* (Fig. 4A; Supplemental Table S3). *At5g35935* is adjacent to the promoter region of *At5g35940* which was down-regulated in both *rrp6l1-2* and *ahl10-1*. ChIP assays found low ψ_w_-induced enrichment of AHL10 at several AT-rich sites in the region between *At5g35935* and *At5g35940* (Fig. 4C). This stress-induced association was not observed for a nearby region in the *At5g35940* gene body or a distal off-site (EIF4a promoter). A similar enrichment pattern was observed for AHL10^S314A^, consistent with the idea that the phosphonull mutation in the C-terminal region of AHL10 does not interfere with the DNA binding region in the N-terminal part of the protein. We also conducted ChIP assays for AT-rich promoter regions within 1 kb of the transcriptional start site for several other genes having decreased expression in *ahl10-1* and *rrp6l1-2* (Supplemental Fig. S8). However, we did not find enrichment of AHL10 on those sites. Possibly AHL10 is associated with more distal sites which influence expression of those genes by formation of chromatin loops which bring AHL10 (and RRP6L1) in closer proximity to the gene or its newly transcribed RNA.

Two studies have previously identified regions of reduced methylation in *rrp6l1-2* (Zhang et al., 2014; Wendte et al., 2017) or conducted transcriptome analysis of *rrp6l1-2* (Zhang et al., 2014). However, we found little overlap between *rrp6l1-2* differentially expressed genes identified by Zhang et al. (2014) and those identified in our analysis (Supplemental Fig. S9A) or between *rrp6l* DEGs identified in any study and *rrp6l1*-related demethylated regions (Supplemental Fig. S9B). The differences in the results could be influenced by slight age difference in seedlings between experiments (11 days in our case, 12-15 days for the other studies). However, the very limited overlap in differentially expressed genes and demethylated regions between these data sets suggests a strong effect of environmental conditions on RRP6L1 action in transcriptional regulation.

## Discussion

Our results demonstrate that RRP6L1 is involved in low ψ_w_ response, that AHL10 and RRP6L1 interact and, that AHL10 S314 phosphorylation may facilitate AHL10 function as an auxiliary factor that promotes RRP6L1 chromatin association. Both RRP6L1 and AHL10 protein levels were high in meristem tissue indicating that this is where AHL10- RRP6L1 interaction mainly occurs and where AHL10 and RRP6L1 exert their effects on growth during low ψ_w_ stress. High levels of AHL10 and RRP6L1 in meristem can explain their effects on expression of meristem-specific genes such as *STM* and the auxin-related genes such as *WES1* and *DFL1*. In the context of previous work, our results indicate that control of AHL10 phosphorylation by competing action of MPKs versus Clade A PP2Cs (particularly HAI1) in meristems is a new connection of stress-related phospho-signaling to gene regulation and growth suppression during low ψ_w_ stress.

While our experiments focused on the effect of AHL10 phosphorylation and did not seek to investigate the mechanism of RRP6L1-mediated gene regulation, it is of interest to put our results in the context of previous characterization of RRP6L1. The lack of RRP6L1 chromatin recruitment in *ahl10-1* during low ψ_w_ stress is surprising as it indicates that AHL10 has a controlling influence on RRP6L1 compared with other RRP6L1-interacting proteins such as the C-Terminal Domain (CTD) of NRPE1, the largest subunit of RNA Polymerease V (Wendte et al., 2017). In that study the authors speculated that RRP6L1 interaction with the CTD is transient or involves only a small portion of Pol V. Our data suggest that the converse may also be true: CTD interaction may involve only a small portion of RRP6L1, particularly during low ψ_w_ stress. Alternatively, RRP6L1 interaction with AHL10 could precede, or be required for, CTD interaction (or other interactions) that position RRP6L1 at specific chromatin locations. Zhang et al. (2014) noted that it is not known what factors determine RRP6L1 preference for certain sites over others. Determining whether AHL10 binding is competitive with CTD binding, or whether AHL10 binding can influence the 3’-5’ exonuclease activity of RRP6L1 (Zhang et al., 2014) will be questions of interest for future studies. It should also be kept in mind that previous studies of RRP6L1 (Lange et al., 2008; Shin and Chekanova, 2014; Zhang et al., 2014; Ye et al., 2016; Wendte et al., 2017) have only observed RRP6L1 function under unstressed conditions and mainly focused on RRP6L1 effects on non-coding RNAs. Thus, it is unclear how their proposed mechanisms of RRP6L1 function may be altered by low ψ_w_ stress and whether RRP6L1 affects protein coding genes via distinct mechanisms compared to its effects on non-coding RNAs.

The lack of RRP6L1 chromatin recruitment in *ahl10-1* is also surprising given that one may have expected a greater degree of redundancy among the closely related Clade B AHLs. Also surprising is our observation that AHL10 and AHL13 are not genetically redundant, despite their overall high similarity and conserved C-terminal phosphorylation site (Wong et al., 2019; Rayapuram et al., 2021). Instead, *ahl13-1* blocked the *ahl10-1* mutant phenotype of increased growth maintenance during low ψ_w_ stress. While they are not redundant, AHL10 and AHL13 have common regulatory targets. For example, both affect the regulation of JA-related genes (*AOC1, AOS*) in response to low ψ_w_ (for *ahl10-1*) or pathogen infection (for *ahl13-1;* Wong et al., 2019; Rayapuram et al., 2021). These seemingly contradictory results can be interpreted in light of previous evidence that AHLs act as trimers (Zhao et al., 2013). It is presumably these trimeric complexes that recruit other transcriptional regulators; albeit, this hypothesis awaits direct testing *in planta*. AHL13 may act as an anchor for different AHL complexes, thus explaining why it is genetically downstream of AHL10. In this case, the combination of AHL13 with AHL10 gives specificity of recruiting additional regulators such as RRP6L1. Knocking out one of these genes disrupts the composition of AHL complexes and thus disrupts recruitment of interacting proteins and downstream phenotypes in ways more complex than simple redundancy among AHLs.

## Data Availability

RNAseq data has been deposited to Gene Expression Omnibus and is available under accession number GSE101197. All other data supporting the findings of this study are in the paper and supplemental material.

## Supplemental Data

Supplemental Figure S1: AHL10 interactors.

Supplemental Figure S2: *In-vitro* pull down experiments confirm direct interaction of AHL10 and RRP6L1.

Supplemental Figure S3: Complementation of *ahl10-1* with AHL10 phosphonull and phosphomimic mutants under control of the *AHL10* native promoter.

Supplemental Figure S4: AHL10 accumulates in root and shoot meristems.

Supplemental Figure S5: Effect of phospho-null and phospho-mimic mutations on AHL10 sub-nuclear localization pattern.

Supplemental Figure S6: Effect of *RRP6L1, AHL10* and *AHL13* mutants on growth during moderate severity low water potential stress.

Supplemental Figure S7: Gene expression changes in *rrp6l1-2* and *ahl10-1* compared to low water potential stress-induced changes in wild type.

Supplemental Figure S8: ChIP assay of AT-rich promoter sites for other genes with decreased expression in *ahl10-1* and *rrp6l1-2* under low water potential stress.

Supplemental Figure S9: Comparison of our *rrp6l1-2* transcriptome data to genes in or near hypomethylated regions in previously reported analyses.

Supplemental Table S1: Genes up or down regulated in *rrp6l1-2* compared to Col-0 wild type in the unstressed control treatment.

Supplemental Table S2: Gene ontology analysis of genes up or down regulated in *rrp6l1-2* in the unstressed control treatment.

Supplemental Table S3: Genes up or down regulated in *rrp6l1-2* compared to Col-0 wild type in the low water potential stress treatment (−0.7 MPa, 96 h).

Supplemental Table S4: Primers used for cloning, genotyping, mutagenesis, gene expression analysis and ChIP.

## Acknowledgements

We thank David Favero (Riken) for providing cDNA clones of AHL27 and AHL29; Ming-Che Shih (Agricultural Biotechnology Research Center, Academia Sinica) for providing the *35S:GFP* transgenic line; Toshisangba Longkumer for experimental assistance, Ji-Ying Huang and Mei-Jane Fang (Live Cell Imaging core facility, Institute of Plant and Microbial Biology) for microscopy assistance; Wen-Dar Lin (Bioinformatics Core Facility, Institute of Plant and Microbial Biology) for assistance with transcriptome data analysis and Shih-Shan Huang for laboratory assistance. This research was supported by an Academia Sinica Investigator Award (AS-IA-108-L04) and a National Science and Technology Council of Taiwan grant (MoST 110-2311-B-001 -019) to P.E.V.

## Author Contributions

P.E.V. conceived research, M.M.W., Y.-C.B. and P.E.V. designed experiments, M.M.W., X.- J..H and Y.-C.B. performed experiments, M.M.W., X.-J..H, Y.-C.B. and P.E.V. analyzed data, P.E.V. wrote the manuscript with assistance from M.M.W. All authors read and approved the manuscript.

