## Supplemental Figures S1-S9 for "AHL10 phosphorylation determines RRP6L1 chromatin association and growth suppression during water stress"

**A**

| Accession number | Gene Name | X-Gal assay | Frequency |
|---|---|---|---|
| AT4G12080 | AHL1 | strong | 10 |
| AT4G25320 | AHL3 | strong | 4 |
| AT2G33620 | AHL10 | strong | 7 |
| AT4G17950 | AHL13 | strong | 4 |
| AT1G54440 | RRP6L1 | strong | 1 |
| AT4G33260 | ATCDC20.2, CDC20.2, CELL DIVISION CYCLE 20.2 | intermediate | 1 |
| AT4G05320 | POLYUBIQUITIN 10, UBI10, UBIQUITIN 10, UBQ10 | weak | 2 |
| AT5G10390 | H3.1, HISTONE 3.1, HTR13 | weak | 1 |
| AT5G54900 | ATRBP45A, RBP45A, RNA-BINDING PROTEIN 45A | weak | 1 |
| AT2G04880 | WRKY1, ZAP1, ZINC-DEPENDENT ACTIVATOR PROTEIN-1 | weak | 1 |
| AT5G04320 | ATSGO2, SGO2, SHUGOSHIN 2 | weak | 1 |
| AT1G51690 | PROTEIN PHOSPHATASE 2A REGULATORY SUBUNIT B ALPHA ISOFORM | weak | 1 |
| AT2G45980 | ATG8-INTERACTING PROTEIN 1, ATI1 | weak | 1 |
| AT3G50500 | SnRK2.2 | weak | 1 |

**B**

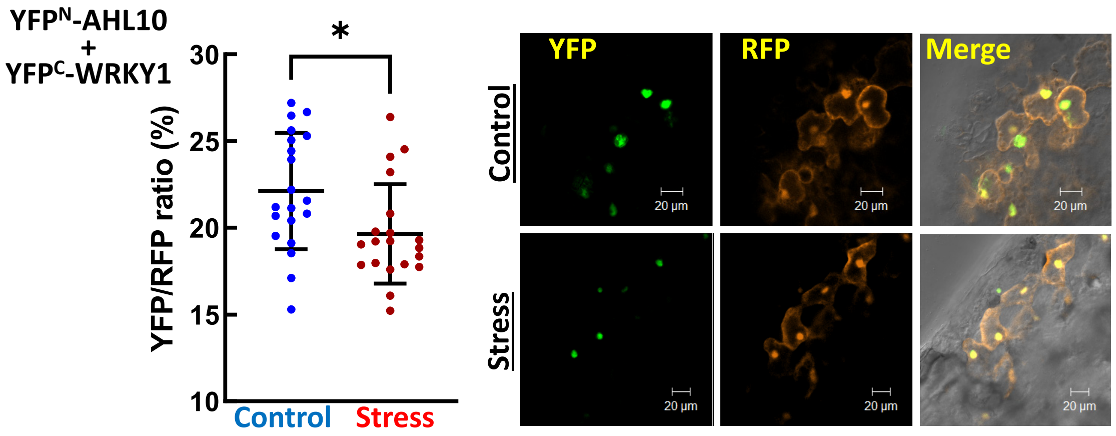


**Supplemental Figure S1: AHL10 interactors (Supports Fig. 1)**

A. Putative AHL10 interactors identified by yeast two hybrid screening. Only proteins with predicted localization in nucleus based on experimental evidence or SUBA prediction (greater than 0.5) are shown. Results of X-gal staining were categorized as weak, intermediate, or strong to indicate the relative strength of the interaction. Frequency refers to the number of independent clones of a particular gene found in the yeast two hybrid screen.

B. rBiFC test of AHL10 interaction with WRKY1 under control and stress (-1.2 MPa) conditions. Quantitation of YFP/RFP ratio from images collected in two independent experiments, along with representative images of the control and stress treatments, are shown.

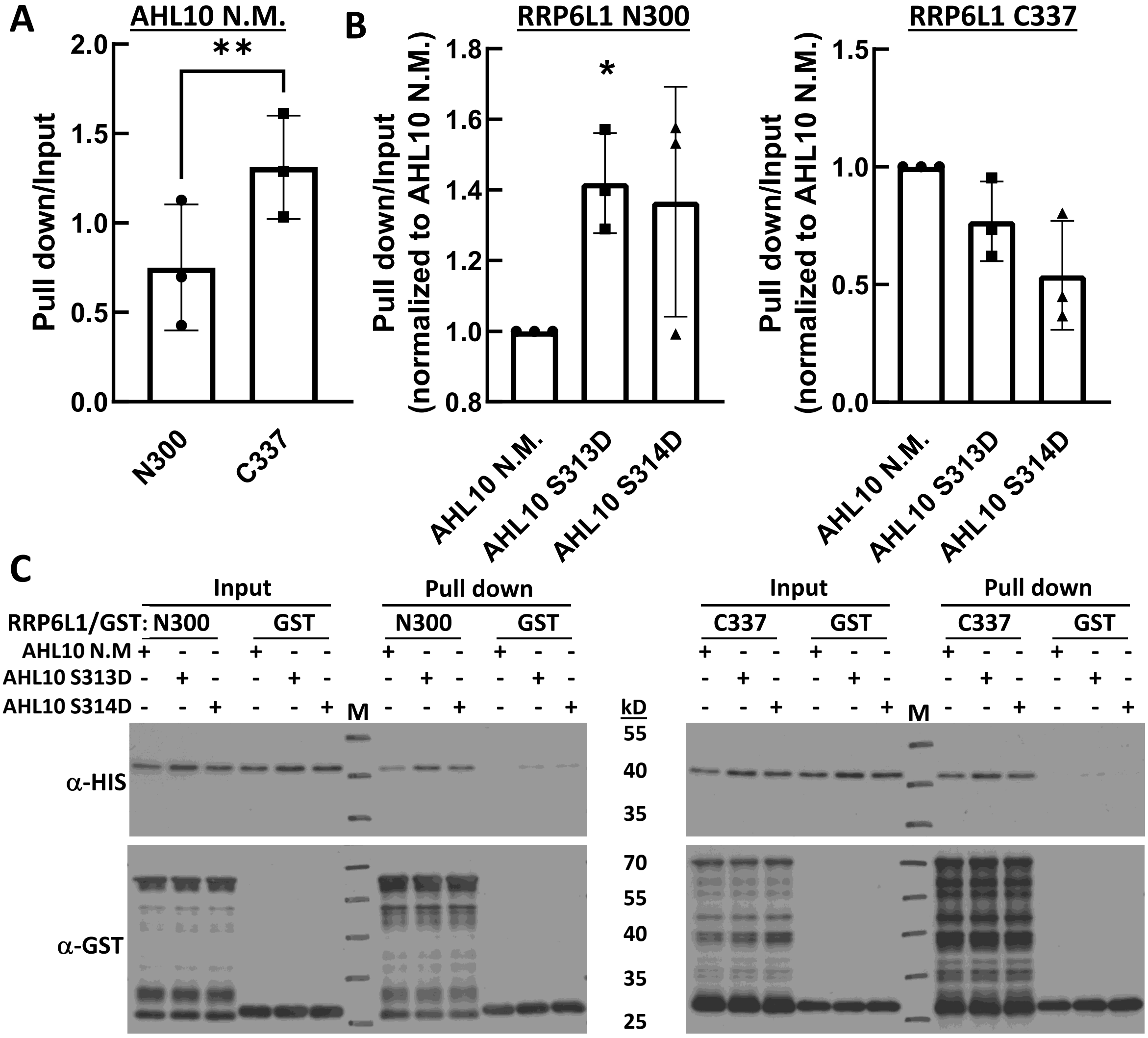


**Supplemental Figure S2: *In-vitro* pull down experiments confirm direct interaction of AHL10 and RRP6L1. (Supports Fig. 1)**

A. Quantitation of relative band intensities of AHL10 (non-mutated AHL10, AHL10 N.M.) pull-down with either the N-terminal 300 amino acids (N300) or C-terminal 337 amino acids (C337) of RRP6L1. GST-tagged RRP6L1 fragments (or free GST) were used to capture HIS-tagged AHL10. Note that we were unable to express full length RRP6L1 in E. coli for pull down assays, consistent with previous reports. Intensity of the pull down band was quantified relative to the input band for three replicate assays.
B. Comparison of RRP6L1 N- and C-terminal interaction with wild type AHL10 (AHL10 N.M.) or phosphomimic versions of AHL10 (AHL10 S313D and S314D). Intensity of the pull down bands of the phosphomimic AHL10 was quantified relative to the band intensity of wild type AHL10 for three replicate assays.
C. Representative immunoblots from one of the experiments quantified in A and B. Error bars indicate S.D. and asterisks (*) indicate significant differences based on T-test.

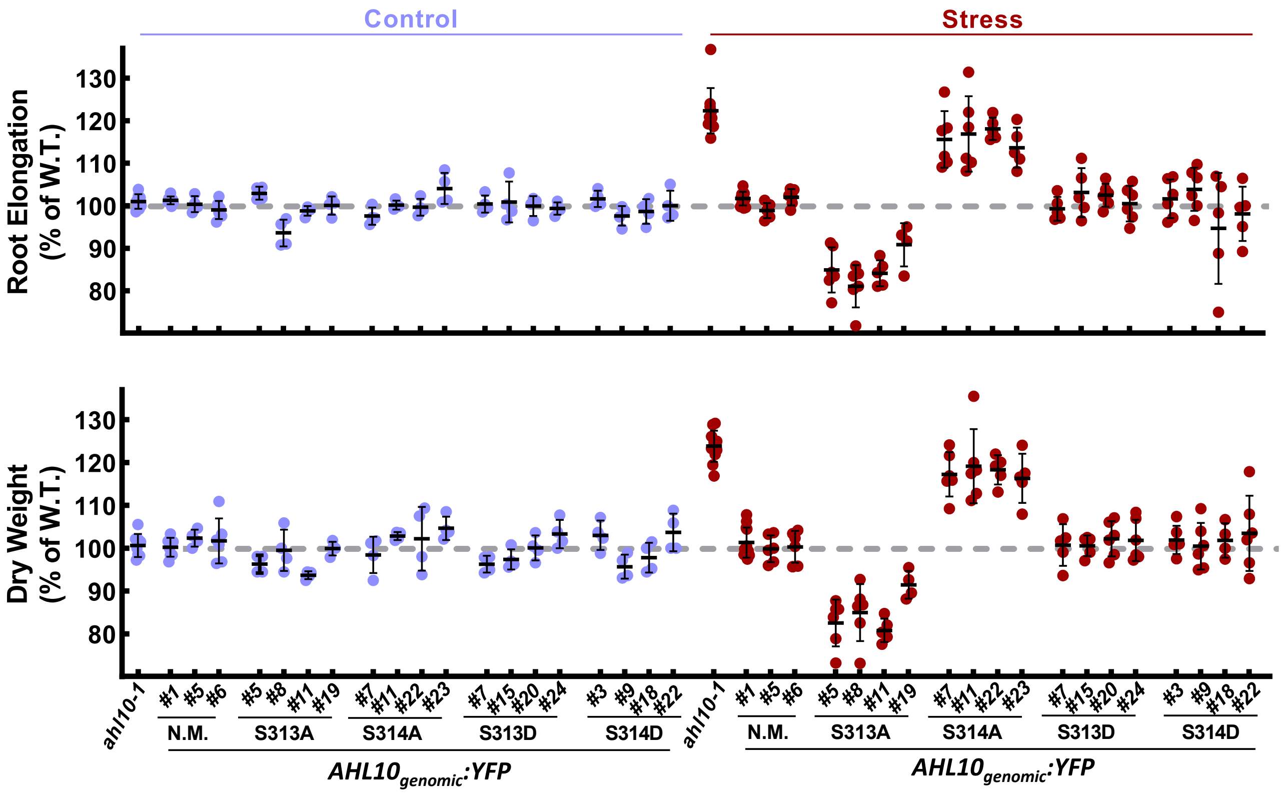


**Supplemental Figure S3: Complementation *of ahl10-1* with AHL10 phosphonull and phosphomimic mutants under control of the *AHL10* native promoter. (Supports Fig. 1 and 2)**

The genomic fragment containing AHL10 (promoter and gene body up to the stop codon) was fused to YFP (C-terminal fusion). The same construct was produced for AHL10 phosphonull (S313A, S314A) and phosphomimic (S313D, S314D) mutants. Growth assays on PEG-agar plates were conducted using homozygous T3 lines of each construct under control and stress (-0.7 MPa) conditions. Data points shown are from individual agar plates where the primary root elongation and seedling dry weight are expressed relative to wild type growing on the same plate. Data are from 2 independent experiments (2 or 3 plates quantified per experiment) are shown. Error bars indicate the standard deviation.

Consistent with previous results obtained with *35S:YFP-AHL10* lines (Wong et al., 2019), the wild type *AHL10* construct (labeled as N.M. for "Non-Mutated") fully complemented the *ahl10-1* phenotype of increased growth maintenance at low water potential. Also consistent with previous results, the S313A version of AHL10 was hyperactive and suppressed growth below the wild type level (presumably because the S313A mutation led to more phosphorylation of S314) while the S314A version of AHL10 failed to complement the mutant phenotype. Both the S313D and S314D versions of AHL10 fully complemented ahl10-1. None of the lines had altered growth in the unstressed control. The data confirmed that S314 phosphorylation is required for AHL10 function in suppressing growth at low water potential.

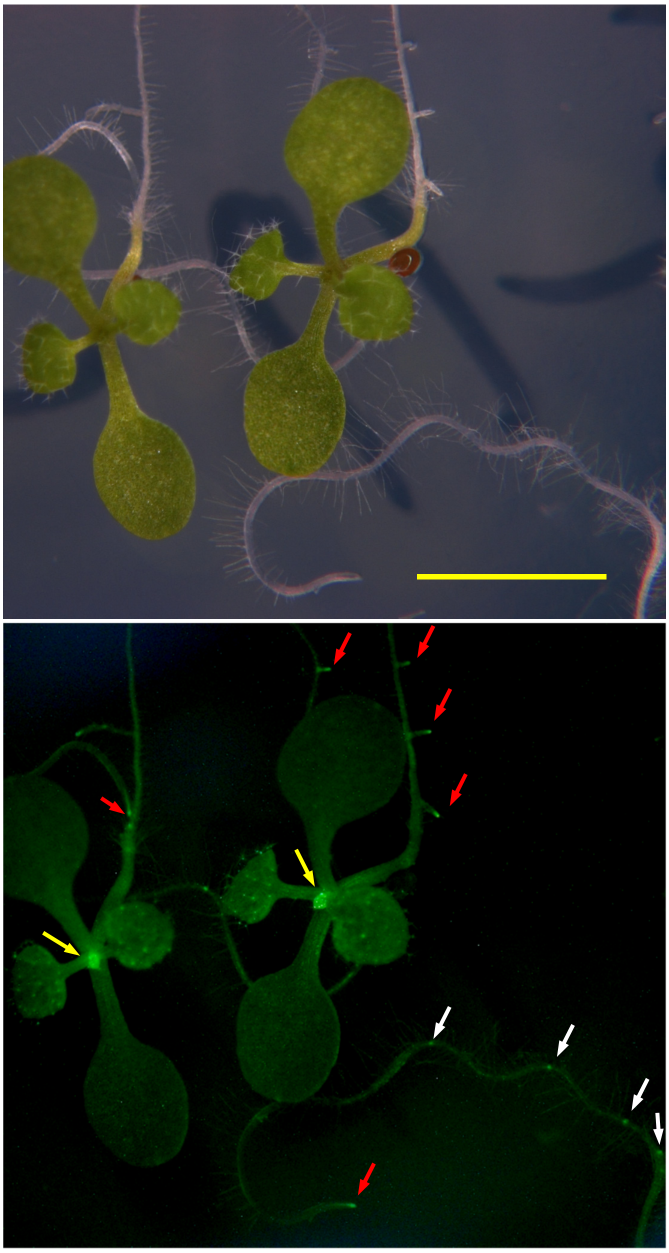

**Supplemental Figure S4: AHL10 accumulates in root and shoot meristems (Supports Fig. 1).**
Representative image of *AHL10genomic:YFP* expressing transgenic plants. Images are from 7 day-old plants in the unstressed control treatment. Images were collected with a fluorescence microscope (Zeiss Axio Imager Z1) to access the pattern of AHL10-YFP protein accumulation at the whole plant level. Note high levels of AHL10-YFP in shoot meristem (yellow arrows) and root meristems (red arrows). AHL10-YFP accumulation also seemed to mark the position of lateral root emergence along the primary root (white arrows). Scale bar indicates 5 mm.

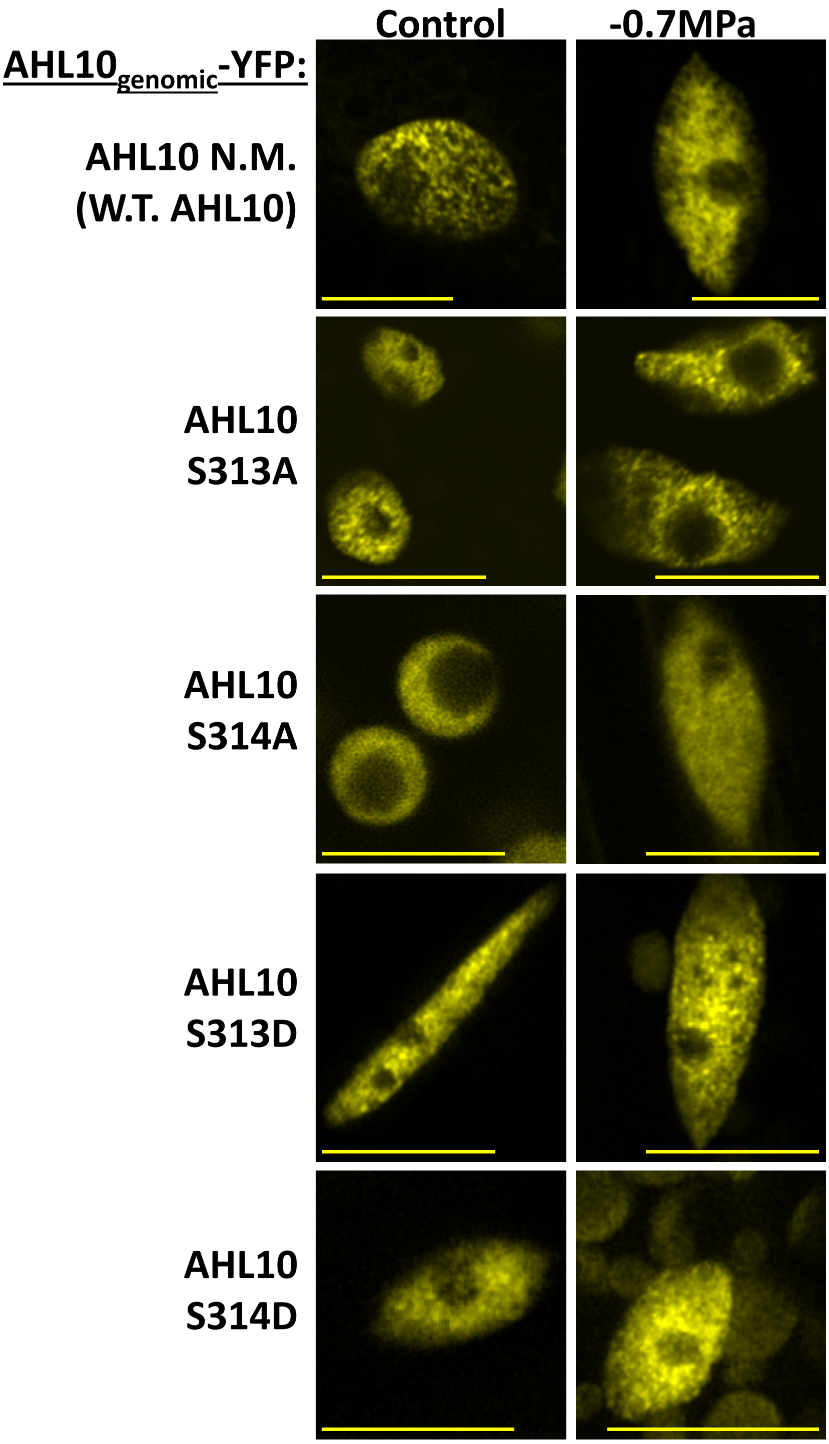


**Supplemental Figure S5: Effect of phospho-null and phospho-mimic mutations on AHL10 sub-nuclear localization pattern. (Supports Fig. 1 and 2).**
Nuclei were imaged from root cells of transgenic lines transformed with the genomic fragment of AHL10 (native promoter and gene body up to the stop codon) fused to YFP. Nuclei were imaged from seedlings under unstressed control conditions or after low water potential treatment (-0.7 MPa for 96 h). Scale bars indicate 10 µm.

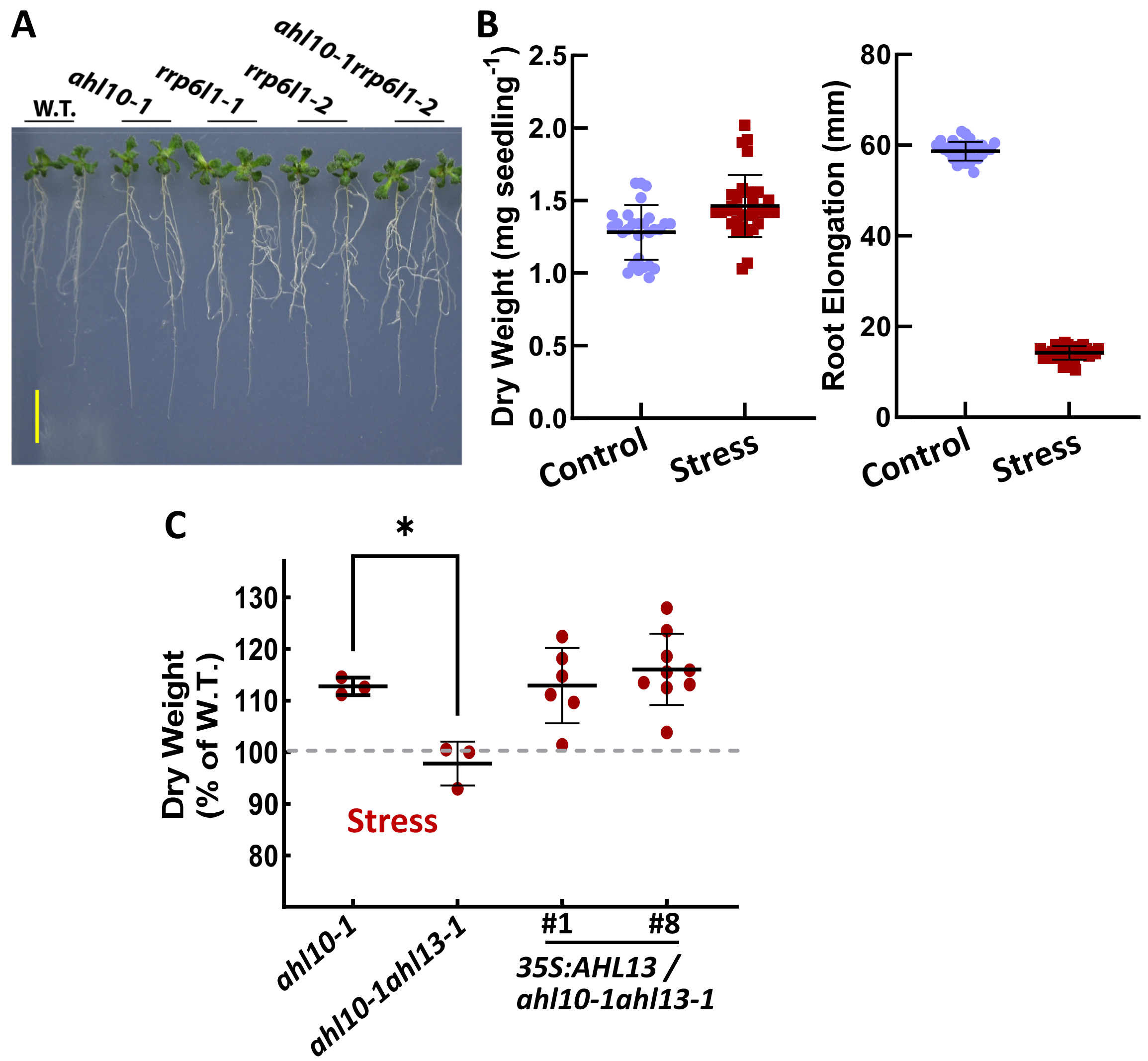


**Supplemental Figure S6: Effect of RRP6L1, AHL10 and AHL13 mutants on growth during moderate severity low water potential stress (Supports Fig. 3).**

A. Representative seedlings in the low water stress treatment from the experiments shown in Fig 3A.

B. Wild type seedling dry weight and root elongation used to normalize the mutant data shown in Fig. 3A. Data are combined from three independent experiments.

C. Complementation of *ahl10-1ahl13-1* with *35S:AHL13* restores the increased growth maintenance seen in *ahl10-1* during low water potential stress. Assay conditions were the same as described for Fig. 3A. Data are means ± S.D. from data from three independent biological experiments. Asterisk (*) indicates significant difference compared to *ahl10-1* (ANOVA, $P \leq 0.05$).

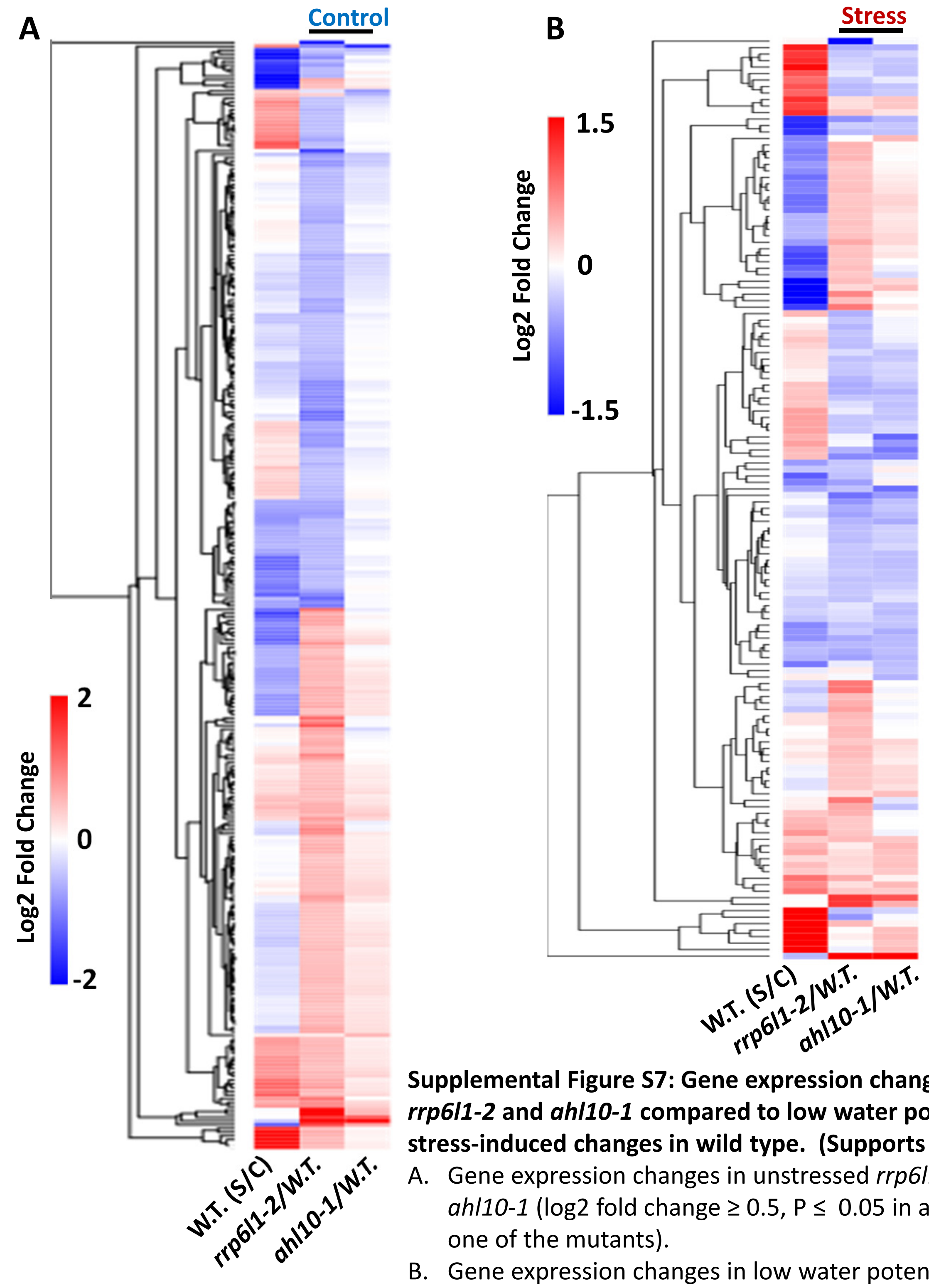


**Supplemental Figure S7: Gene expression changes in *rrp6l1-2* and *ahl10-1* compared to low water potential stress-induced changes in wild type. (Supports Fig. 4)**

A. Gene expression changes in unstressed *rrp6l1-2* or *ahl10-1* (log2 fold change ≥ 0.5, P ≤ 0.05 in at least one of the mutants).
B. Gene expression changes in low water potential stress-treated *rrp6l1-2* or *ahl10-1* using a slightly relaxed cutoff for log2 fold change (≥ 0.3) than the heat map shown in Fig. 4A.

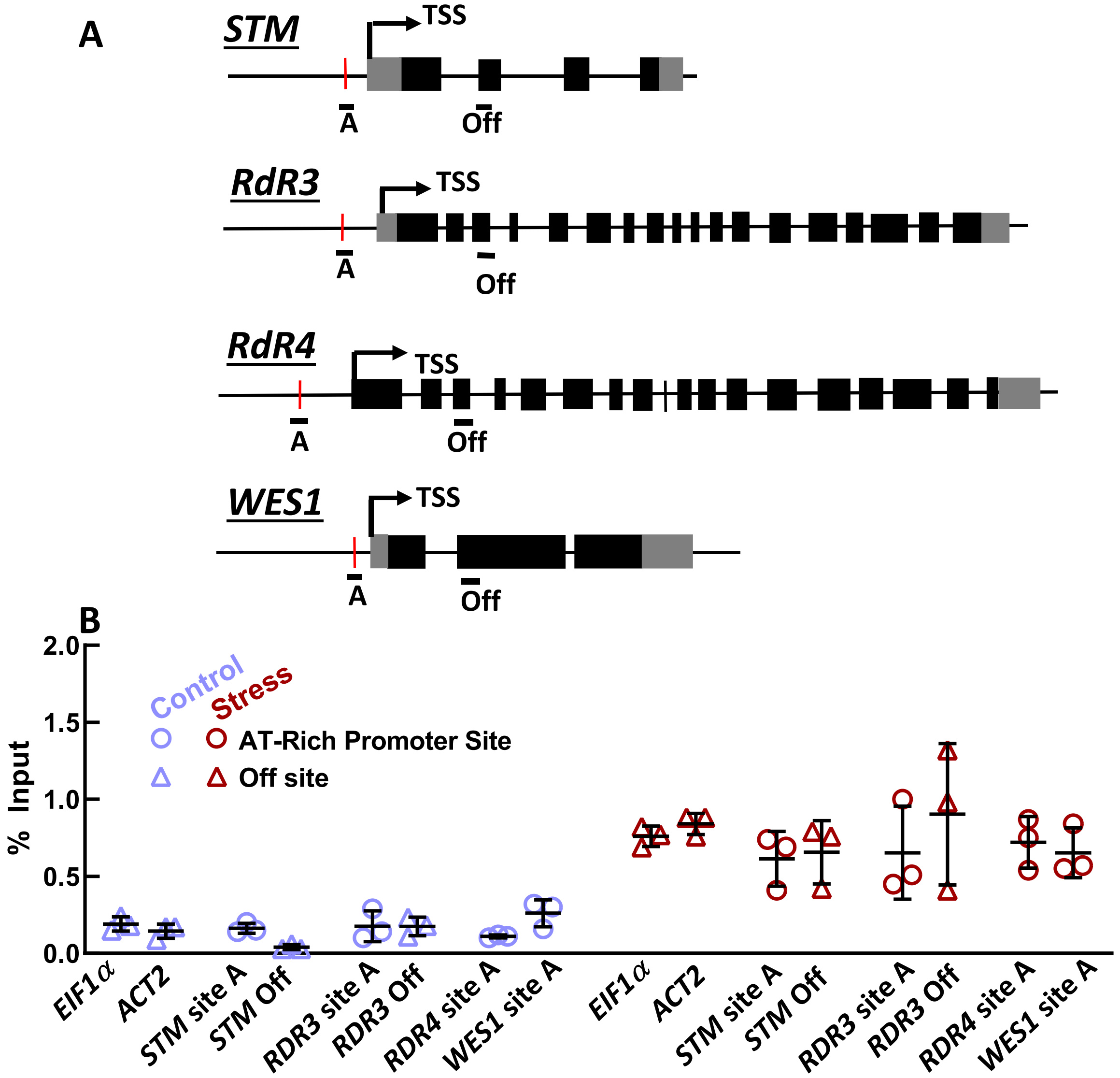


**Supplemental Figure S8: ChIP assay of AT-rich promoter sites for other genes with decreased expression in *ahl10-1* and *rrp6l1-2* under low water potential stress. (Supports Fig. 4)**

A. Gene diagrams showing the location of AT-rich promoter sites and off sites analyzed by ChIP assay.
B. ChIP assays results. *EIF1α* and *ACT2* promoter sites were used as negative controls. Data are from three biological replicates (error bars indicate the S.D.).

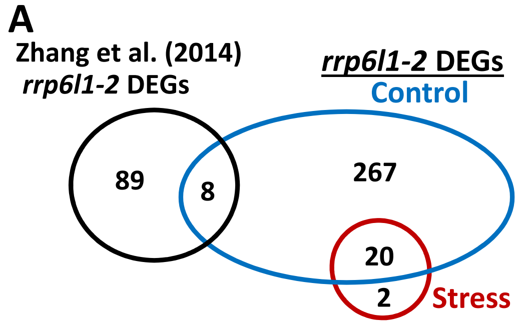


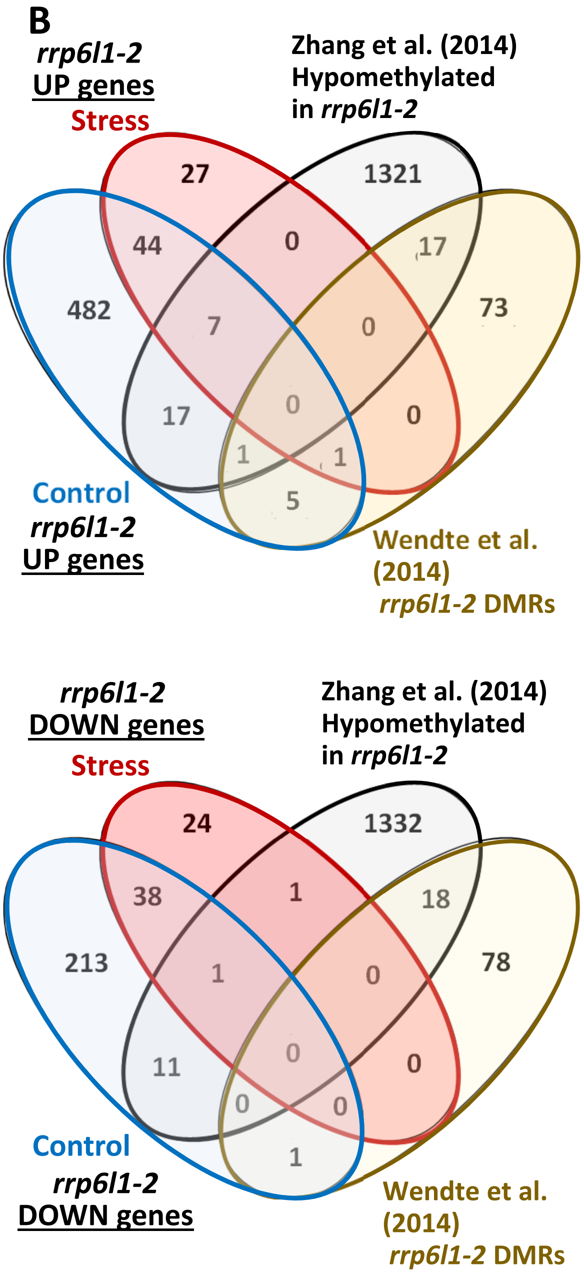


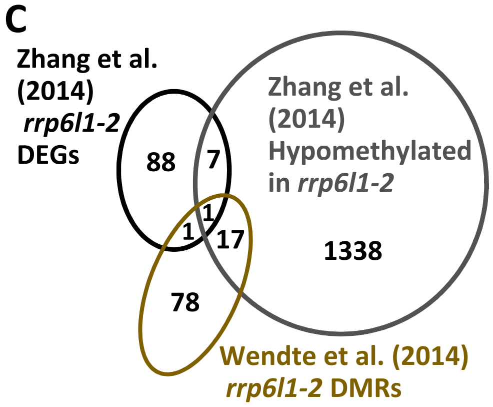


**Supplemental Figure S9: Comparison of our *rrp6l1-2* transcriptome data to genes in or near hypomethylated regions in previously reported analyses (Supports Fig. 4).**

A. Comparison of *rrp6l1-2* differentially expressed genes (DEGs) under stress or control treatments in our experiments (log2 fold-change ≥ 0.5, adjusted P ≤ 0.05) versus differentially expressed genes reported by Zhang et al. (2014) using a 2-fold expression change cutoff.

B. Expression of differentially expressed genes identified in our analysis of *rrp6l1-2* to genes adjacent to regions found to by hypomethylated in rrp6l1 mutant compared to wild type in genome wide methylation analysis conducted by Zhang et al. (2014) and Wendte et al. (2017). The definition of a gene associated with a hypomethylated region (1 base pair or more overlap between the hypomethylated region and the gene) was used to extract a list of genes from the list of Demethylated Regions (DMRs) reported in Wendte et al. (2017).

C. Comparison of differentially expressed genes in *rrp6l1-2* reported by Zhang et al., (2014) to genes overlapping or adjacent to hypomethylated regions reported by Zhang et al. (2014) or Wendte et al. (2017).
